# Turanose induced WOX5 restores symbiosis in the *Medicago truncatula* cytokinin perception mutant *cre1*

**DOI:** 10.1101/830661

**Authors:** Anindya Kundu, Firoz Molla, Maitrayee DasGupta

## Abstract

Rhizobia-legume interaction recruits cytokinin-signaling that causes local auxin accumulation for the induction of nodule primordia in the cortex. Since sugar signaling can trigger auxin responses and regulate developmental processes, we explored whether sugar treatments could rescue cre1. Here we demonstrate that turanose, a non-metabolizable sucrose analogue can recover functional symbiosis in cytokinin perception mutant *cre1*. Additionally, turanose significantly upregulated the expression of WUSCHEL-related homeobox 5 (*MtWOX5*) which prompted us to check if ectopic expression of WOX5 could rescue *cre1*. Overexpression of WOX5 from *Arachis hypogaea (AhWOX5*), but not the intrinsic *MtWOX5* could completely restore functional symbiosis in *cre1* though both WOX5 (Mt and Ah) were functionally equivalent in inducing the expression of cytokinin inducible transcription factor Nodule Inception (*NIN*). Among the tested markers for cytokinin and auxin responses, significant differences were noted in the expression of IAA-Ala Resistant3 (*MtIAR33*), an auxin conjugate hydrolase. Turanose and *AhWOX5* overexpression resulted in upregulation of *MtIAR33* that further increased significantly in presence of rhizobia. On the other hand, *MtIAR33* expression was unaffected in *MtWOX5* overexpressed roots suggesting deconjugation driven auxin pool to be critical for rescuing symbiosis in *cre1*. We hypothesize a working model for sugar and WOX5 mediated rescue of symbiosis in *cre1*.

**One sentence summary:** Activation of sugar-WOX5 signaling axis restores root nodule symbiosis in cytokinin perception mutant *cre1*

## INTRODUCTION

In root nodule symbiosis (RNS) rhizobia-legume interaction activates the SYM pathway which in turn activates cytokinin response for the induction of a *de novo* meristem in the root cortical cells (Frugier et al., 2008). Cytokinin-signaling causes local auxin accumulation at the site of incipient nodule primordia by modulating the expression of auxin transporters (Plet et al., 2011; Ng et al., 2015). These phytohormonal signals and the SYM pathway together reprogram the cortical cells and regulate their division, ultimately building a nodule primordium for the endocytic accommodation of the symbionts (Mathesius et al., 2000; Suzaki et al., 2012).

Several evidences projected the hierarchical organization of the Nod factor-dependent activation of SYM-Pathway and cytokinin signaling pathway (Frugier et al., 2008). For example, Nod factor induced accumulation of cytokinin by upregulation of several cytokinin biosynthesis genes indicates its precedence to cytokinin signaling(Held et al., 2014; van Zeijl et al., 2015). This explains why the majority of Nod factor-induced transcriptional changes were absent in cytokinin perception mutant *cre1* in *M. truncatula*(van Zeijl et al., 2015). Cytokinin signaling through CRE1 (CYTOKININ RESPONSE 1) is required for the induction of NIN, a key transcription factor required for nodule organogenesis as well as for infection (Gonzalez-Rizzo et al., 2006; Plet et al., 2011; Vernié et al., 2015; Liu et al., 2019). Cortical activation of cytokinin signaling that is prerequisite for nodule organogenesis is also dependent on NIN (Vernié et al., 2015). Genetic evidences further confirm this hierarchy, when constitutive activation of CCaMK (*snf1*) fails to trigger spontaneous nodulation in absence of cytokinin signaling in *cre1* whereas constitutive activation of cytokinin receptor LHK1 (*snf2*) triggered nodulation in *dmi2* (Madsen et al., 2010). Cytokinin oxidase/dehydrogenase expressed during nodule development restricts the level of active cytokinin for balancing its positive role in nodule organogenesis with its negative effect on rhizobial infection at the root epidermis (Reid et al., 2016). Therefore, apart from being the key endogenous signal for nodule development and differentiation (Plet et al., 2011; Kundu and DasGupta, 2018), cytokinin also maintains the homeostasis of the symbiotic interaction indicating that the *SYM* pathway and cytokinin signaling are connected through feedback loops (Miri et al., 2016).

Hierarchically auxin is projected to act next to cytokinin during nodule development (Plet et al., 2011; Suzaki et al., 2012). Several evidences indicated directional transport and local accumulation of auxin to be important for the development of nodule primordia and accordingly application of auxin transport inhibitors is sufficient to induce nodule-like cell division (Rightmyer and Long, 2011; Ng et al., 2015). Accumulation of auxin could be either due to regulation of its transport or biosynthesis and cytokinin is known to regulate both these processes (van Noorden et al., 2006; Pernisová et al., 2009; Jones et al., 2010). The response regulators (RRs), functioning downstream to cytokinin receptors modulate auxin signaling via SHY2 mediated downregulation of auxin transporters (Ioio et al., 2008; Moubayidin et al., 2010). Recent evidence revealed that cytokinin controls flavonoid concentration in roots and local application of flavonoid could recover nodulation in the cytokinin perception mutant *cre1* of *M. truncatula* (Ng et al., 2015). Though auxin accumulation by its biosynthesis depends on cytokinin inducible NIN expression (Schiessl et al., 2019), the auxin transport inhibitors can cause pseudonodule formation in mutants like *nsp2* and *nin* suggesting auxin accumulation to be driven by diverse upstream cues (Rightmyer and Long, 2011). Overall auxin accumulation and response appear to be the final call for initiation of cortical cell division during RNS (Ng et al., 2015).

Sugars play a role in plant development and several evidences indicate a crosstalk between sugar signaling and auxin responses. For example, (i) there are mutants that are commonly resistant to both sugar and auxin responses (Ohto et al., 2006), (ii) auxin biosynthetic genes are modulated by sugar status (LeClere et al., 2010; Sairanen et al., 2012) and (iii) there is a connection between PAT and sugar metabolism (Stokes et al., 2013). Moreover, sugar signaling through auxin-based signal transduction is also responsible for specific developmental responses like trehalose mediated inhibition of root elongation and glucose affecting root architecture (Wingler et al., 2000; Mishra et al., 2009). Sugar signaling is also connected with auxin response of the conserved WUS (WUSCHEL)/WOX (WUSCHEL-related homeobox gene)-CLV regulatory system that are involved in meristem maintenance. For example, WOX9 is typically required for auxin response in Shoot Apical meristem (SAM) and sucrose can compensate for loss of WOX9 (*stip*), that arrests growth soon after germination (Wu et al., 2005). Also, the expression of WOX5, which maintains localized auxin maxima in the root apical meristem (RAM) is induced by auxin and turanose, a non-metabolizable sucrose analogue (Gonzali et al., 2005).

CRE1, an orthologue of *AtAHK4*, is a membrane-bound cytokinin receptor necessary for nodulation in *M. truncatula* (Gonzalez-Rizzo et al., 2006; Plet et al., 2011). Earlier reports indicated that the *M. truncatula cre1* mutant is defective in auxin accumulation in the root cortex following *S. meliloti* inoculation (Ng et al., 2015). A plausible hypothesis could be that sugar signaling would rescue cytokinin perception mutant *cre1* by inducing auxin responses. Sucrose is an important metabolite transported to nodules and sucrose transporters are upregulated with rhizobial infection but their specific role in signaling during nodule development is still oblivious (Lalonde et al., 2004; Ayre, 2011). The dual role of sucrose as an energy source and a signaling molecule is evidenced by many experiments and turanose, a non-metabolizable analogue of sucrose provides an opportunity to study its role as a signaling molecule (Sinha et al., 2002). Herein we show that turanose treatment upregulates both cytokinin and auxin responses and successfully rescues the nodulation efficiency in *cre1*. Turanose treatment induced the expression of WOX5, and ectopic expression of *WOX5* can also restore symbiosis in *cre1*.

## RESULTS

### Turanose restores root nodule symbiosis in *M. truncatula* cytokinin perception mutant *cre1*

Based on several evidences that highlight the crosstalk between sugar signaling and auxin responses we wanted to explore whether sugar treatments could rescue rhizobial symbiosis in cytokinin perception mutant *cre1* (LeClere et al., 2010; Ljung et al., 2015). Since sugars can act as energy source, as well as signaling molecules, we disentangled these functions by using sucrose as well as turanose, a non-metabolizable sucrose which is previously reported to trigger auxin response in *Arabidopsis* (Gonzali et al., 2005). Both *A17* and *cre1* roots were pre-treated with sucrose and turanose (10^−4^M to 10^−2^M) for 1 week before being infected with *Sinorhizobium meliloti Sm2011-pBHR-mRFP*. Under these conditions, there was no significant change in root or shoot length or the lateral root number in either *A17* or *cre1* plants in presence of turanose whereas sucrose strongly affected both root and shoot growth (Supplementary Fig.1). On the other hand, while 10^−3^M turanose treatment could completely recover symbiosis in *cre1*, treatment with sucrose at the same concentration was unable to restore symbiosis (Fig.1; Supplementary Table 1). These findings suggested that sucrose act as a signaling molecule rather than as a metabolizable energy source for the activation of RNS in *cre1*. At 3 weeks after infection (WAI), there were ~1.4 ± 0.9 nodules in untreated *cre1* roots whereas turanose treatment resulted in formation of ~12 ± 3.3 nodules per plant (Fig.1A). In untreated A17 there were ~21 ± 2.9 nodules at 3WAI which increased to ~28 ± 8.7 nodules upon turanose treatment (Fig.1A). Almost 42% of the turanose rescued nodules in *cre1* were pink nodules (Fig.1B and E), with proper colonization and differentiation of bacteroids (Fig.1I). For the rest of the nodules that were white and merged (Fig.1B and F), rhizobia were entrapped in cortical infection threads (Fig.1J). The increase in nodule number in turanose treated A17 was due to significant increase in merged white nodules (Fig.1B and H). In these nonfunctional white nodules rhizobia were entrapped in infection threads in the infection zone as opposed to the proper colonization and differentiation of bacteroids in the pink functional nodules (Fig.1K-L). It may be noted that formation of merged nodules was a common feature in turanose treated *cre1* and A17 roots indicating that turanose induced loss of spatial distribution is independent of CRE1 mediated cytokinin signaling.

**Figure 1:**
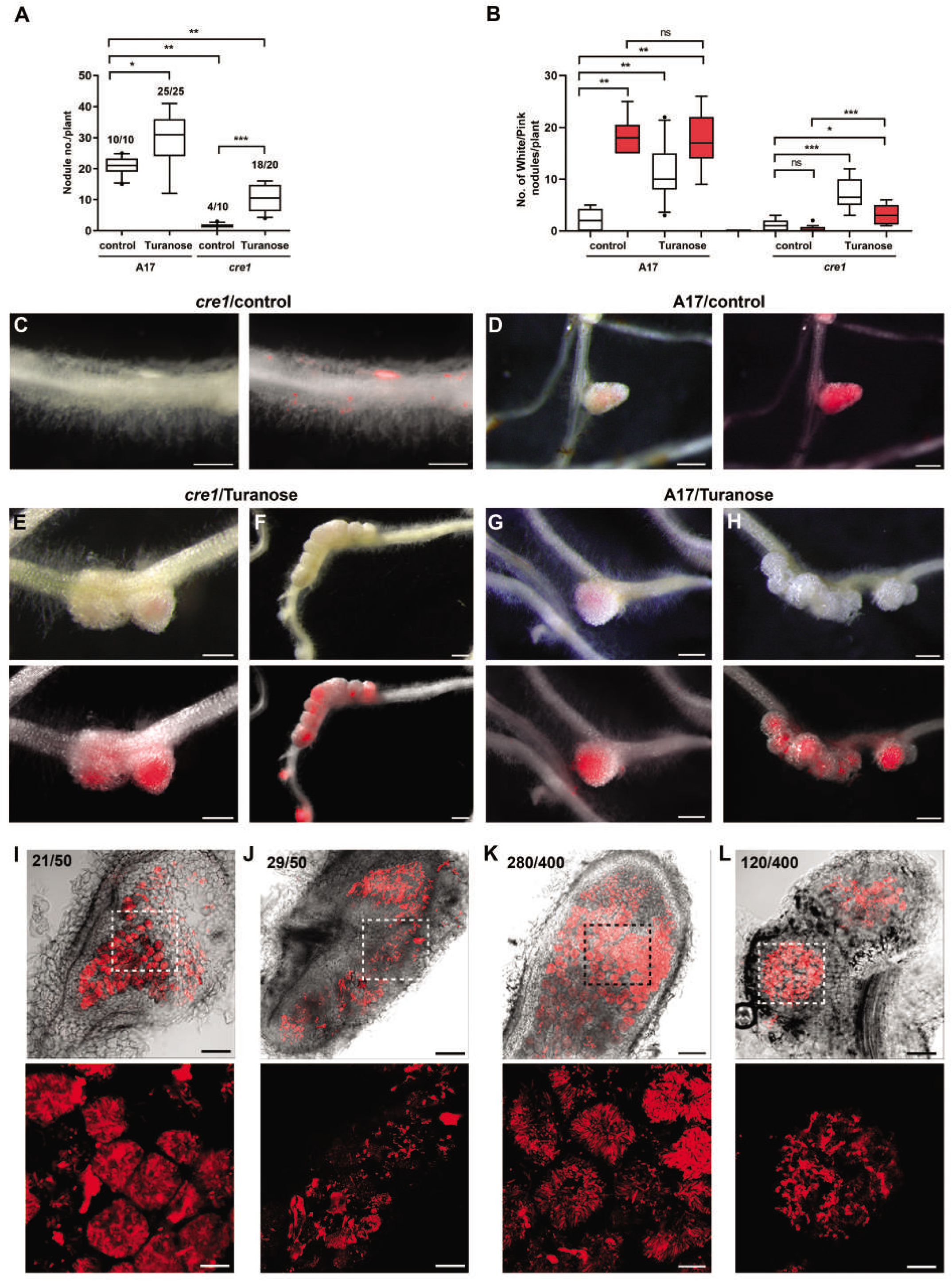
Turanose mediated recovery of symbiosis in *cre1* mutants. **(A-B)** Box plot represents (A) total nodule number per plant and (B) total white (white box) and pink nodules (red box) under control (mock) and turanose treatment (10^−3^M) in A17 and *cre1* after infection with *Sinorhizobiummeliloti Sm2011-pBHR-mRFP*. Number above each box in (A) indicates the number of root systems with nodules per total number of scored roots. Mann-Whitney test were used to assess significant differences, where ***, ** and * indicates P<0.0005, <0.002 and <0.02 respectively. **(C-H)** Morphology of infection event or nodules in *cre1* and A17 3WAI under (C-D) control condition and (E-H) after turanose treatment where (E and G) solitary pink nodules and (F and H) merged white nodules. Representative micrograph of control and turanose treated nodulated roots indicated as bright field and bright field + mRFP merged. **(I-L)** Ultrastructure analysis of nodules in *cre1* and A17 after turanose treatment, where the perforated box represents the infection zone with its corresponding enlarged view below the respective images. Total number of each type of nodules observed out of the total observed nodules are indicated for each panel. (upper panel) Brightfield + mRFP merged and (lower panel) mRFP. Scale bar: C-H= 500μm, I-L (upper panel) = 100μm and (lower panel) = 10μm.

### Turanose mediated distribution of cytokinin and auxin response maxima in transgenic roots of composite *M. truncatula cre1 mutants*

To understand the probable mechanism behind turanose mediated rescue of symbiosis in *cre1* we analyzed the cytokinin and auxin responses at early stages of rhizobial infection. As a readout for cytokinin response we used *pTCS:GUS*, the cytokinin responsive two-component signaling (TCS) reporter that uses the target binding sequence of ARR-Bs (Müller and Sheen, 2008; Zürcher et al., 2013). The *pTCS:GUS* transformed hairy roots of both *A17* and *cre1* were pre-treated with 10^−3^M turanose for 1 week before being infected with *Sinorhizobium meliloti* for 24 hours. Histochemical analysis was done for spatial localization of GUS activity where staining was strictly done for 15 minutes to highlight the noted contrast in presence of turanose. In uninfected mock treated A17, *pTCS:GUS* was only detectable in meristematic zone (MZ) which in infected roots spread into the epidermal layer of the both MZ and root hair zone (RHZ) (Fig.2A-B). Earlier report have also shown a similar activation of *TCSn:GUS* during early stages of rhizobial invasion, with maximal expression in epidermal and outer cortical cells in the nodulation-competent RHZ (Jardinaud et al., 2016). Interestingly in turanose treated A17 roots even in absence of rhizobia, the expression of *pTCS:GUS* was noted in the epidermal layers of both MZ and RHZ which at 24hpi further extended in the cortex and along the root stele (Fig.2C-D). Such expansion of *pTCS:GUS* in inner cortical cells is usually noted at later states of rhizobial infection (Vernié et al., 2015). Under identical conditions *pTCS:GUS* was not detectable in mock treated *cre1*, but upon infection, it was noted near the outer layers of MZ (Fig.2E-F). On the other hand, turanose treated *cre1* roots have a significantly enhanced *pTCS:GUS* expression in the MZ and in the epidermal and cortical layers of RHZ. Thus turanose appears to have changed the cytokinin signaling landscape in *M. truncatula* roots and the increased expression of *pTCS:GUS* in *cre1* roots indicate CRE1 to spatially restrict the level of cytokinin responses (Fig.2G-H).

**Figure 2:**
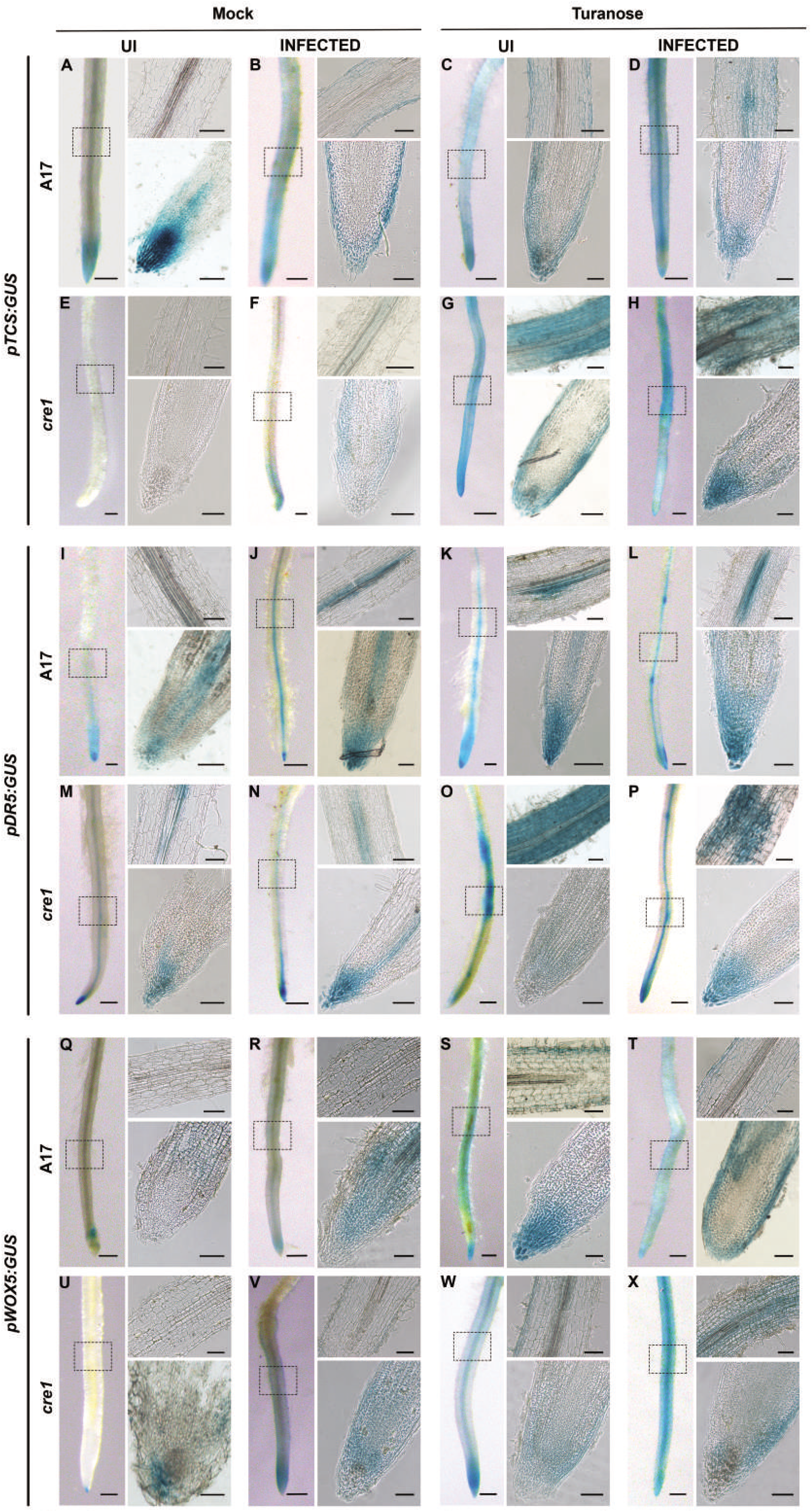
Effect of turanose treatment on *pTCS:GUS, pDR5:GUS* and *pWOX5:GUS* expression pattern in A17 and *cre1* roots. **(A-H)** *pTCS:GUS* expression in (A-D) A17 and (E-H) *cre1* roots,(I-P) *pDR5:GUS* expression in (I-L) A17 and (M-P) *cre1* roots and (Q-X) *pWOX5:GUS* expression in (Q-T) A17 and (U-X) *cre1* roots under following conditions. **(A-H)** mock treated uninfected (A, E), infected (B, F), turanose treated uninfected (C, G) and infected (D, H). **(I-P)** mock treated uninfected (I, M), infected (J, N), turanose treated uninfected (K, O) and infected (L, P). **(Q-X)** mock treated uninfected (Q, U), infected(R, V), turanose treated uninfected (S, W) and infected (T, X). All plants were mock or Turanose (10^−3^M) treated for 7 days followed by infection with *Sinorhizobium meliloti Sm2011-pBHR-mRFP*for 24 hours before being harvested for GUS staining. Every root image has its corresponding longitudinal section of the root tip and the root hair zone (box). At least 5 independent roots were analyzed for each condition from three independent experiments. Scale bars = 300 μm (left panel single root), Scale bar = 50μm (right panel upper and lower longitudinal section).

As a readout for auxin response we used *pDR5:GUS*, that uses multiple tandem repeats of a highly active synthetic auxin response element (Ulmasov et al., 1997; Sabatini et al., 1999). Under identical conditions, in wild type A17 roots *pDR5:GUS* was noted along the stele near the MZ but not in the RHZ (Fig.2I). This expression increased along the root stele in the RHZ in infected A17 roots but activity in the root cortex was very low (Fig.2J). Earlier observations also reported auxin response to be restricted to vascular bundles and pericycle in *M. truncatula* roots (van Noorden et al., 2007; Demina et al., 2019). Interestingly in turanose treated roots this increased auxin response along the root stele in RHZ was noted in absence of rhizobia and this pattern remained unchanged in turanose treated infected roots (Fig.2K-L). In uninfected *cre1* roots the pattern of auxin response was similar to A17 except that it was also detected in the RHZ along the stele and the pattern remained similar in the infected roots as well (Fig.2M-N). However in turanose treated *cre1*, *pDR5:GUS* expression was significantly higher in RHZ with GUS staining in the epidermal and cortical layers (Fig. 2O). Concomitantly, 1μM IAA treatment of *M. truncatula* was previously reported to cause a similar auxin response that spread along the whole root system (van Noorden et al., 2007). Thus, our observations suggest a similar surge of auxin inside the root upon turanose treatment. Such patches of *pDR5:GUS* expression in *cre1* were also noted in the cortex of turanose treated infected roots but in a lesser frequency (Fig.2P). It is well documented that cytokinin signaling through CRE1 is required for the accumulation of auxin in cortical cells to initiate nodule organogenesis (Ng et al., 2015). Since auxin response is significantly high in turanose treated *cre1* roots, our results additionally demonstrate that CRE1 has a significant role in restricting the level and spatial spread of the auxin response. In summary, turanose treatment induced significant cytokinin and auxin responses in the RHZ of the *cre1* roots and thus indicating a probable cause behind the rescue of nodulation phenotype.

### Turanose treatment elevated the expression of *MtWOX5* in *cre1*

The crosstalk of WOX family of homeobox transcription factors with the sugar and phytohormonal signaling networks play a key role in meristem development and maintenance (Wu et al., 2005; Wahl et al., 2010; Kong et al., 2016). Earlier observations indicated *MtWOX5* expression to coincide with the site of nodule primordium formation where the first divisions are initiated in the pericycle, endodermis, and inner cortical cells (Osipova et al., 2011). We were curious to check how the changes in phytohormonal responses in turanose treated roots affected the expression of *MtWOX5* in the early stages of rhizobial infection (Fig.2A-P). Experimental conditions were therefore kept unchanged and *MtWOX5* expression was followed using *pWOX5:GUS* in turanose treated roots in presence (24hpi) and in absence of rhizobia. qRT-PCR estimation revealed *MtWOX5* expression to be elevated in presence of turanose with the effect being more pronounced in *cre1* as compared to A17. This increased expression of *MtWOX5* in *cre1* indicates cytokinin signaling through MtCRE1 to have a role in restricting *MtWOX5* expression in A17 (Supplementary fig.2).

We then analyzed the spatial expression pattern of *pWOX5:GUS* and compared it with cytokinin (*pTCS:GUS*) and auxin (*pDR5:GUS*) responses (Fig.2A-P). In absence of turanose in uninfected roots of both A17 and *cre1*, expression of *pWOX5:GUS* was restricted to root apex (Fig.2Q and U). Upon infection *pWOX5: GUS* spread in the inner layers of the MZ in A17, whereas in *cre1* roots it spread in both outer and inner layers. This indicates that CRE1 has a role in restricting the expression of *MtWOX5* (Fig.2R and V). In turanose treated A17 roots, both in presence or in absence of rhizobia, *pWOX5:GUS* expression expanded to the epidermal layers of RHZ. This highlights an overlap of *pWOX5:GUS* with cytokinin response pattern and suggests a role of cytokinin signaling in turanose mediated transcriptional activation of *MtWOX5* (Fig.2C, S and T). In uninfected roots of turanose treated *cre1*, *pWOX5:GUS* expression was also noted in epidermal layers of RHZ indicating that CRE1 may not have a role in turanose mediated *WOX5* expression (Fig.2W). In turanose treated infected *cre1* roots *pWOX5:GUS* expression was primarily centered towards stele showing an overlap between auxin response, but was also present throughout the cortex that shows overlap with both cytokinin and auxin responses (Fig.2H, P and X). Thus, both cytokinin and auxin responses show significant overlap with *MtWOX5* expression and could be involved together in determining the level and distribution of *MtWOX5* in turanose treated infected roots of *cre1*. Intriguingly, previous reports also indicated *MtWOX5* expression to be a function of both cytokinin and auxin response during the onset of nodule development (Osipova et al., 2012).

### Ectopic expression of WOX5 rescues nodulation in *cre1*

Earlier observations have demonstrated that *STIP (WOX9*) overexpression (OE) restores SAM in cytokinin sensing mutants(Wu et al., 2005). The significant increase in *MtWOX5* expression in turanose treated *cre1* roots raises the question whether OE-WOX5 would rescue the symbiotic phenotypes in *cre1*. Phylogenetically WOX5 from determinate nodule forming legumes like *Arachis hypogaea, Glycine max, Vigna angularis, Cajanus cajan* and *Phaseolus vulgaris* and indeterminate nodulators like *Cicer arietinum, Pisum sativum, Medicago truncatula*and *Trifolium subterraneum* distinctly clusters in a distance tree and therefore could be amenable to distinct regulations (Supplementary fig.3). While acropetal auxin transport inhibition is essential for indeterminate nodule development, it is dispensable for determinate nodulation (Ng and Mathesius, 2018). We therefore attempted to rescue *cre1* by OE-*MtWOX5* as well as OE-*AhWOX5* from a determinate legume like *Arachis hypogaea. Mt*WOX5 is 86% similar to *Ah*WOX5 and the homeobox domain is highly conserved with 96% similarity (Supplementary fig.3).

Full length *MtWOX5* and *AhWOX5* was amplified from cDNA prepared from respective plant roots. *Mt*WOX5 codes for a protein of 184aa (XP_003616581.1) and *Ah*WOX5 codes for a protein of 215aa (KT820790). Transformed roots of *cre1* overexpressing *MtWOX5* and *AhWOX5* were inoculated with *S. meliloti Sm2011-pBHR-mRFP* or *Sm1021-pXLGD4-lacZ*. Infection threads (ITs) were detected within 2WAI in both empty vector transformed and OE-*MtWOX5 cre1* roots (Fig.3A, Supplementary Fig.4). But in both the cases IT development was abnormal (A) where ~75% ITs were stalled in epidermal cortical barrier and ~20% were stalled in microcolonies in root hair. In both cases, cortical cell divisions were noted but these primordia-like structures remain uninfected (Supplementary Fig.4C and F). These problems in progress of infection were previously reported for *cre1* (Plet et al., 2011), which indicates that OE-*MtWOX5* could not overcome the infection related abnormalities. On the other hand, within 2WAI ~75% of the ITs in OE-*AhWOX5* roots were normal (N) (Fig.3A). The normal ITs progressed toward the subtending nodule primordia generated in the cortex to ensure proper rhizobial accommodation (Supplementary fig.4G-I). It is worth noting that with increased incidences of normal ITs in OE-*AhWOX5* roots, there was a significant reduction in the total number of ITs observed in comparison to empty vector or OE-*MtWOX5* roots (Fig.3B). In accordance with the abnormal progression of ITs there was no improvement of nodulation efficiency in OE-*MtWOX5 cre1* roots even at 6WAI (n=10) (Fig.3C; Supplementary Table 2). But overexpression of *AhWOX5* led to significant nodulation in *cre1* roots by 4WAI where the total number of nodules was comparable to wild type A17 roots (Supplementary fig.4; Supplementary Table 2). Almost 75% of these nodules were pink (Fig.3D) and could reduce acetylene (Fig. 3G) with ultrastructure analysis revealing properly differentiated bacteroids (Fig.3M). Rest of the nodules that were whitish, resembled the nodules formed in *cre1* or OE-*MtWOX5 cre1* roots, where bacteria were trapped within a network of infection threads in the infection zone (Fig.3H-M). The level of expression of *MtWOX5* and *AhWOX5* were comparable in respective OE-*cre1* roots indicating an additional layer of regulation required for MtWOX5 to become functional in *cre1* (Fig.3E and F). Since overexpression of neither *AhWOX5* nor *MtWOX5* had any effect in nodule number or their organization in A17, it is clear that their signaling output is under the homeostatic control of a large network of fate governing factors (Supplementary Table 2).

**Figure 3:**
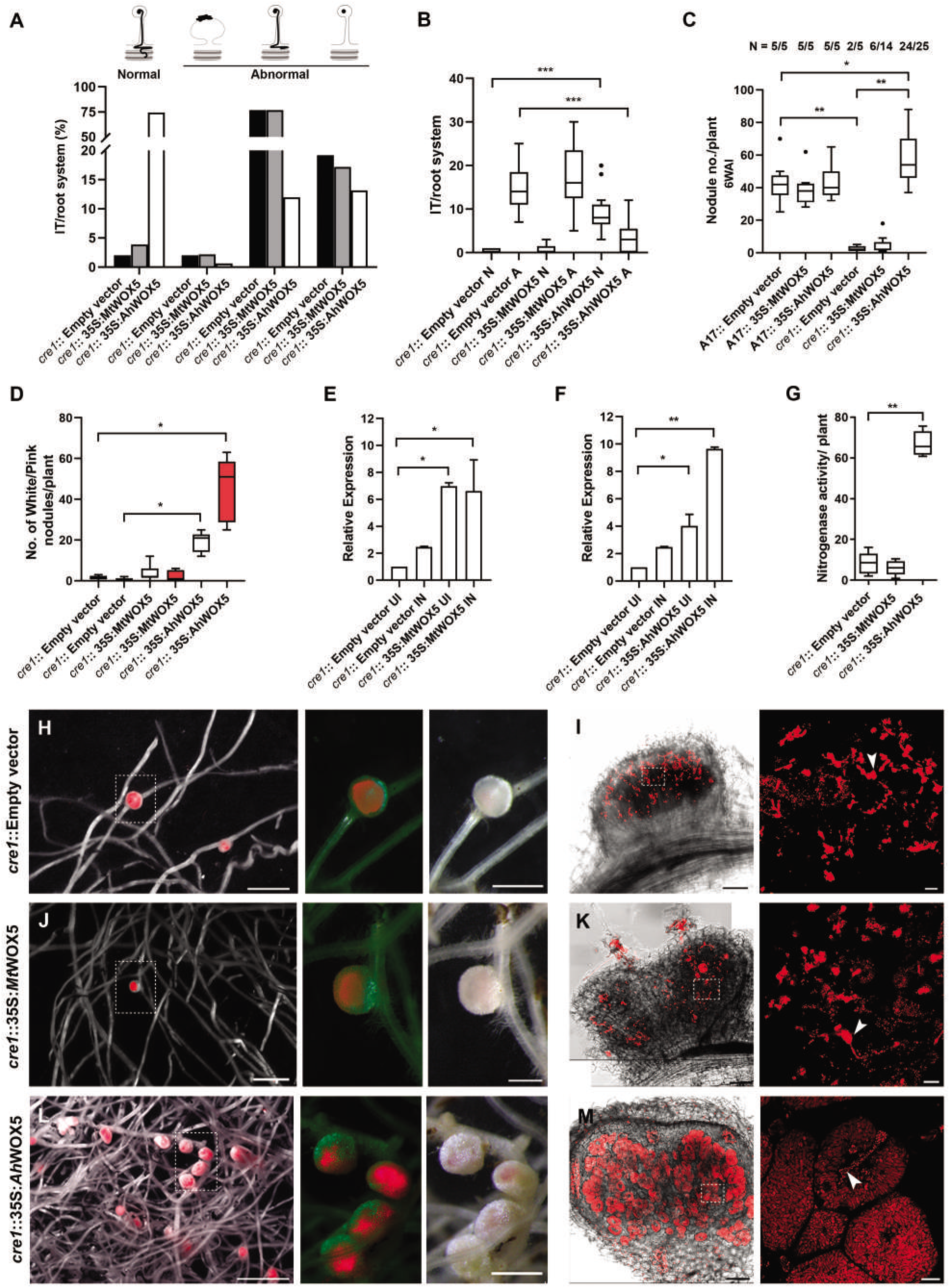
Rescue of symbiosis in *cre1* by ectopic expression of *AhWOX5*. **(A)** Percentile projection of epidermal infection thread (IT) observed 2WAI with *Sm1021-pXLGD4-lacZ* in *cre1* roots transformed with empty vector, *p35S::eGFP-MtWOX5* and *p35S::eGFP-AhWOX5*. Each of the infection events has been roughly classified into 4 categories: (i) successful cortical invasion and intracellular colonization (ii) Stalled in nodule apex, (iii) stalled in epidermal cortical barrier and (iv) stalled in microcolonies in root hair. Category (i) is considered normal whereas Category (ii-iv) is considered as abnormality.**(B)** Box plot representing cumulative number of normal and abnormal ITs. **(C-D)** Box plot represents (C) total nodule number per plant and (D) total white (white box) and pink nodules (red box) per plant in empty vector, *p35S::eGFP-MtWOX5* and *p35S::eGFP-AhWOX5* transformed hairy-root systems in A17 and *cre1* respectively, harvested 6WAI with *Sinorhizobiummeliloti Sm2011-pBHR-mRFP*. n indicates the number of root systems with nodules per total number of scored roots. Mann-Whitney test was used to assess significant differences, where ***, ** and * indicates P<0.0005, 0.002 and <0.02 respectively.**(E-F)** qRT-PCR analysis of (E)*MtWOX5* and (F) *AhWOX5* relative to empty vector transformed (*cre1*) roots normalized against *MtActin*. **(G)** Acetylene reduction assay (nmole C2H4/h/mg nodule) in indicated root system. (E-G) Histogram represent an average of three biological replicates each having n>4 plants and error bar represents SD. Mann-Whitney test was used to assess significant differences, where ***, ** and * indicates P<0.0005, 0.002 and <0.02 respectively. **(H-M)** Morphology of nodules in transgenic hairy-roots of *cre1* mutant 4WAI with *Sm2011-pBHR-mRFP* (H, J and L). Representative micrograph of transgenic roots indicated as bright field + mRFP merged and perforated box is represented as eGFP + mRFP merged and bright field. Scale bar = 1mm and box = 500μm. (I, K and M) Ultrastructure of nodules developed in *cre1* mutant transformed with different constructs indicated as bright field+ mRFP merged where the perforated box in the infection zone is enlarged separately. Scale bar: I, K and M = 100μm and box = 10μm. Arrow indicate infection thread.

### *MtWOX5* and *AhWOX5* overexpression has contrasting effects on expression of auxin and cytokinin response markers

To understand how OE-*AhWOX5* roots have altered cytokinin and auxin responses at early stages of infection (24hpi), we checked the expression of several markers as a readout of these phytohormonal responses in OE-*AhWOX5* roots of both A17 and *cre1*. The same markers were also evaluated in OE-*MtWOX5* roots and turanose treated roots under otherwise identical conditions to have possible clues towards understanding the similarities and differences in their ability to restore symbiosis in *cre1*.

For cytokinin response the expression of *MtCRE1* and its paralogous *MtHK2* and *MtHK3* and several type-A RRs such as *MtRR4, MtRR8* and *MtRR9* that are induced during nodule organogenesis were monitored (Gonzalez-Rizzo et al., 2006; Tirichine et al., 2007; den Camp et al., 2011). It was noted earlier that the late and inefficient nodule development in *cre1* is due to the redundant involvement of receptors *MtHK2* and *MtHK3* (Held et al., 2014; Boivin et al., 2016). In both A17 and *cre1* at early stages of infection, OE-*WOX5* (*Mt* or *Ah*), had little or no effect in the level of expression of *MtCRE1, MtHK2* and *MtHK3* (Fig.4A-C). Turanose treatment however had interesting consequences. In A17 roots, turanose downregulated *MtCRE1* but not *MtHK2* and *MtHK3*, suggesting a CRE1 independent cytokinin perception by these paralogues must be functional in presence of turanose (Fig.4A-C). In *cre1* roots turanose doubled *MtHK2* expression suggesting *MtHK2* expression could be important in turanose mediated recovery of symbiosis in *cre1* (Fig. 4B). There were notable differences in the expression of RRs between OE-*MtWOX5* and OE-*AhWOX5*. Both OE-WOX5 (*Mt* and *Ah*) induced the expression of *MtRR4*, *MtRR8* and *MtRR9* in A17 and *cre1* indicating MtCRE1 independent regulation of their expression in presence of WOX5 (Fig.4D-F). Upon infection, expression of all 3 RRs significantly increased in A17 but in *cre1* their expression was differentially affected. For example, the level of *MtRR8* further increased upon infection but *MtRR9* was unaffected (Fig.4E-F). These effects were however similar in both OE-*MtWOX5* and OE-*AhWOX5*. On the other hand, *MtRR4* expression in infected *cre1* roots was downregulated in OE-*MtWOX5* but significantly upregulated in OE-*AhWOX5* (Fig.4D). This contrast in regulation of *MtRR4* expression in OE-*MtWOX5* and OE-*AhWOX5* might be crucial in determining the difference in their ability to rescue *cre1*. Additionally, similar to OE-*AhWOX5* the turanose treated roots also had increased expression of *MtRR4* in both A17 and *cre1* whereas expression of *MtRR8* and *MtRR9* was unaffected (Fig.4D-F). Thus, for rescue of symbiosis turanose could use MtHK2 as a proxy for MtCRE1, whereas WOX5 might directly target the downstream RRs. This possibility is corroborated by an earlier report which shows that WOX proteins can directly regulate the expression of Type-A RR genes (Zhao et al., 2009).

**Figure 4:**
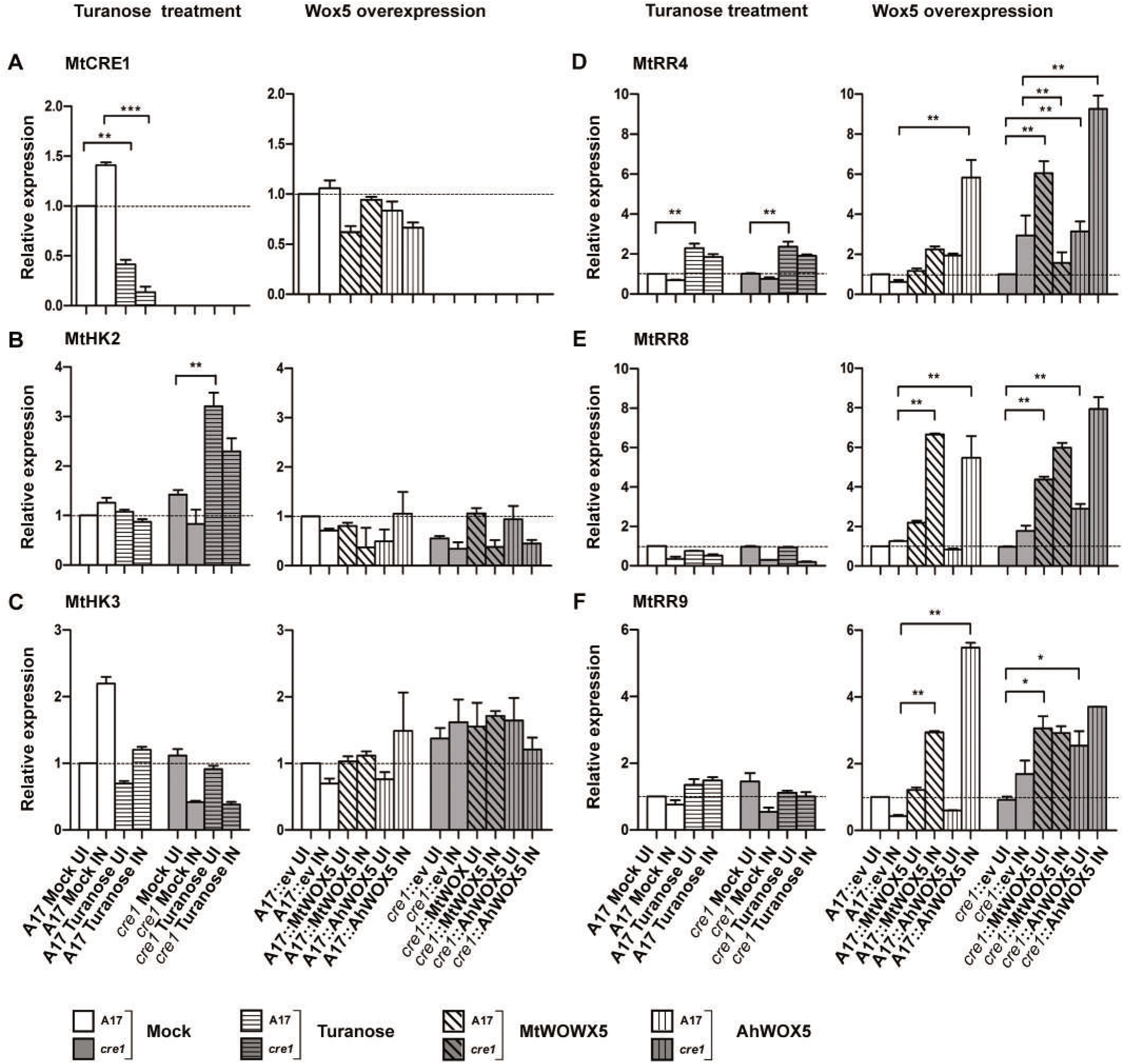
Expression analysis of cytokinin responsive genes under turanose treatment and WOX5 overexpression in A17 and *cre1*. Quantitative reverse transcriptase-PCR analysis of indicated genes in A17 and *cre1* roots **(A, B, C, D, E, F)** treated with turanose (10^−3^M) and water (mock) followed by infection with *Sinorhizobiummeliloti Sm2011-pBHR-mRFP*relative to mock treated uninoculated control or transformed with empty vector, *p35S::AhWOX5* and *p35S::MtWOX5* followed by infection with *Sm2011-pBHR-mRFP* relative to empty vector transformed uninoculated control. *MtActin* was used as a reference gene. Histogram represents an average of three biological replicates each having n>3 plants and error bar represent standard deviation (SD). Mann–Whitney test was used to access significant differences where ***, ** and * represents P < 0.001, <0.01 and <0.05 respectively.

In the early stages of rhizobial infection cytokinin-inducible transcription factor NIN activates LBD16 to promote local accumulation of auxin by triggering its biosynthesis (Schiessl et al., 2019). RNAseq analysis in *cre1* shows that *MtLBD16* and *MtYUCCA2/8* are very feebly expressed in the mutant indicating a role of CRE1 in inducing auxin biosynthesis. We therefore monitored the expression of *MtNIN*, along with *MtLBD16* and *MtYUCCA2/8* as a readout of the auxin response (Fig.5). In both A17 and *cre1, OE-MtWOX5* and OE-*AhWOX5* upregulated the expression of *MtNIN* (~5-10 fold) which further increased with infection (~15-20 fold) (Fig.5A). Also, in both A17 and *cre1*, turanose treatment significantly upregulated *MtNIN* expression in infected roots (~10-25 fold). Thus, till the expression of *MtNIN* the response of *cre1* for OE-*MtWOX5* and OE-*AhWOX5* was identical. As noted earlier, *MtLBD16* expression was very low in *cre1* mutants and turanose has no effect on *MtLBD16* expression in either A17 or *cre1* roots making it unlikely for *MtLBD16* to participate in the Turanose-WOX5 signaling axis (Fig.5B). On the other hand, both OE-*MtWOX5* and OE-*AhWOX5* led to increased expression of *MtLBD16* (~4-5 fold) in *cre1* but not in A17, indicating WOX5 mediated induction of *MtLBD16* to be CRE1 independent and also subject to CRE1 mediated inhibition. Upon infection, only in OE-*AhWOX5* roots there was significant increase in expression of *MtLBD16*. This increase was again significantly more in *cre1* as compared to A17 confirming *AhWOX5* to activate *MtLBD16* in a CRE1 independent pathway (Fig.5B). Similar to *MtLBD16*, turanose treatment had no effect on expression of either *MtYUCCA2* or *MtYUCCA8* indicating that turanose signaling does not directly recruit auxin biosynthesis gene to induce the auxin response (Fig.5C-D). *MtYUCCA2* expression was unaffected by OE-*MtWOX5*, but *MtYUCCA8* expression was upregulated similarly in both OE-*MtWOX5* and OE-*AhWOX5*, and in both A17 and *cre1* indicating a CRE1 independent pathway to activate WOX5 dependent *MtYUCCA8* expression (Fig.5C-D). Local auxin accumulation can also happen by regulating free auxin availability through auxin deconjugation or by affecting auxin efflux transporters (Plet et al., 2011; Deinum et al., 2012). For deconjugation, we chose to monitor the expression of *MtIAR33* as it is evidenced to be highly expressed during nodule organogenesis (Campanella et al., 2008). Unlike in OE-*MtWOX5*, OE-*AhWOX5* resulted in ~10-fold increase in *MtIAR33* expression under uninoculated condition. The *MtIAR33* expression further spiked to ~50-fold post infection in OE-*AhWOX5* in both A17 and *cre1* indicating that its induction under this condition is independent of CRE1 (Fig.5E). Similarly, turanose treatment also resulted in ~150-fold increase in *MtIAR3* expression in infected roots of *cre1* as compared to A17 (~50-fold) suggesting CRE1 might be playing a role in restricting turanose dependent *MtIAR3* expression. Genetic evidence supports the fact that acropetal auxin transport mediated by MtPIN4 plays a very significant role in increasing local auxin concentration during nodule organogenesis (Plet et al., 2011; Ng et al., 2015). However, neither *WOX5* overexpression nor turanose treatment had any effect on *MtPIN4* expression suggesting auxin transport may not be critical in turanose or WOX5 mediated effects (Fig.5F). It is worth mentioning that *MtPIN* expression is not always in concert with the actual auxin transport capacity during nodule development (Plet et al., 2011; Ng et al., 2015).

**Figure 5:**
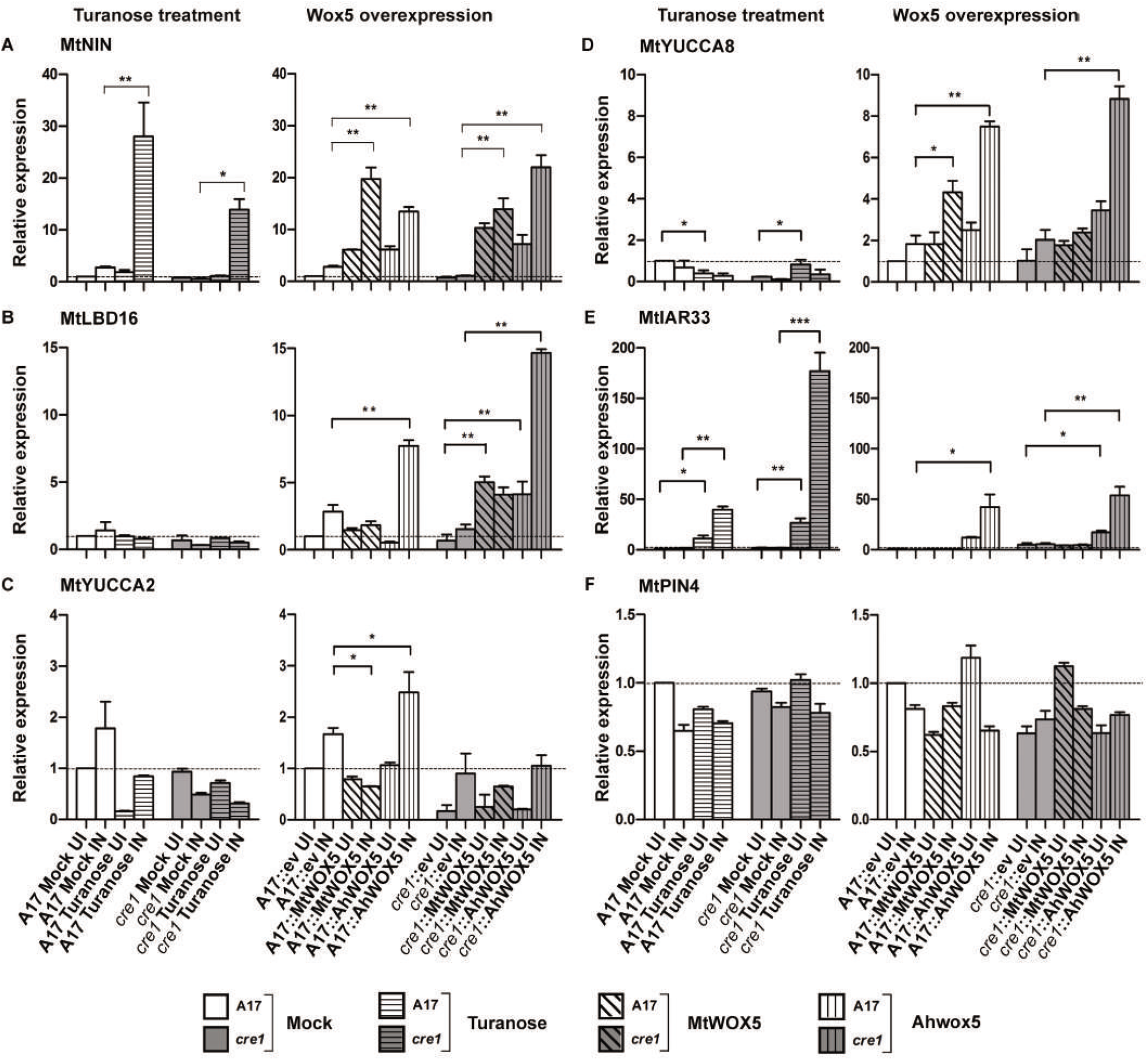
Expression analysis of auxin responsive genes under turanose treatment and WOX5 overexpression in A17 and *cre1*. Quantitative reverse transcriptase-PCR analysis of indicated genes inA17 and *cre1* roots**(A, B, C, D, E, F)** treated with turanose (10^−3^M) and water (mock) followed by infection with *Sinorhizobiummeliloti Sm2011-pBHR-mRFP* relative to mock treated uninoculated control or transformed with empty vector, *p35S::AhWOX5* and *p35S::MtWOX5* followed by infection with *Sm2011-pBHR-mRFP* relative to empty vector transformed uninoculated control. *MtActin* was used as a reference gene. Histogram represents an average of three biological replicates each having n>3 plants and error bar represent standard deviation (SD). Mann-Whitney test was used to access significant differences where ***, ** and * represents P < 0.001, <0.01 and <0.05 respectively.

## DISCUSSION

The symbiotic inefficiency in *cre1* is due to its inability to induce the synthesis of flavonoids that regulate auxin transport for reactivation of cortical cells (Ng et al., 2015). In accordance, treatment with flavonoids allowed auxin transport control and rescued nodulation efficiency in *cre1*. Herein we demonstrate that (i) turanose treatment restores symbiosis in *cre1* (Fig.1). While sugar signaling is known to control several distinct aspects of plant development our results for the first time show its role in RNS. (ii) Turanose induces the expression of *MtWOX5* and ectopic expression of *AhWOX5*, but not *MtWOX5* could restore symbiosis in *cre1* (Fig.2–3). (iii) Both turanose and OE-*AhWOX5* mediated rescue of symbiosis in *cre1* is associated with significant increase in the expression of an auxin conjugate hydrolase *MtIAR33* at early stages of rhizobial infection. As hydrolysis of auxin conjugates are known to be critical in determining free auxin availability (Rampey et al., 2004), we propose this increase in expression of *MtIAR33* downstream to turanose-WOX5 signaling pathway to be responsible for the restoration of symbiosis in cre1 (Fig. 5). Based on our observations we propose a working model for turanose and WOX5 mediated rescue of symbiosis in *cre1* (Fig.6).

**Figure 6:**
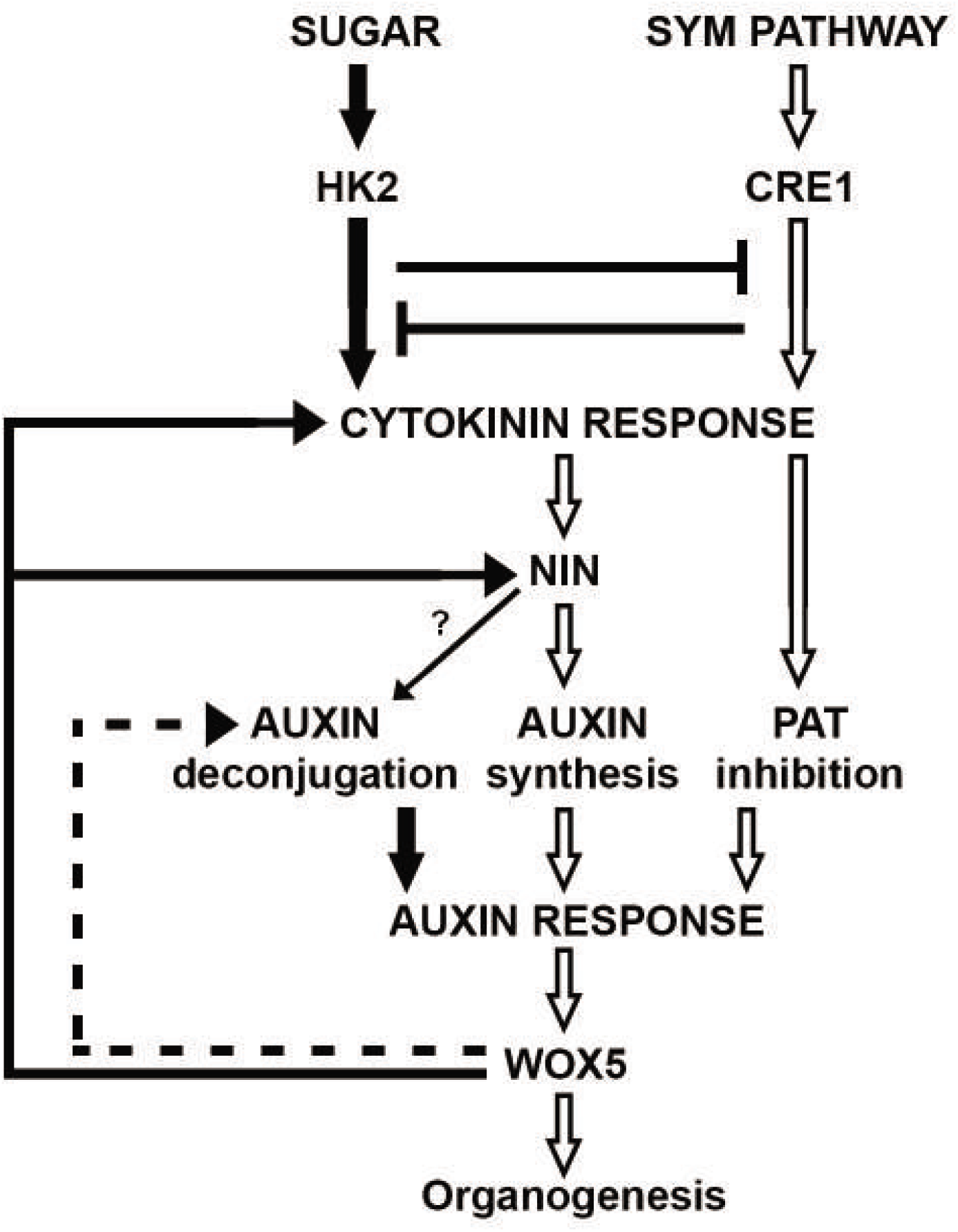
Proposed Model for the role of sugar signaling and *WOX5* expression in restoration of symbiotic efficiency in *M*. truncatula cytokinin perception mutant *cre1*. Open arrows indicate SYM pathway dependent *CRE1* activation that leads to auxin accumulation by auxin biosynthesis and/or auxin transport inhibition. The local auxin accumulation induces *WOX5* to generate the nodule primordia. On the basis of our observations we propose the black arrows indicating sugar induced cytokinin response, *NIN* expression, deconjugation mediated auxin response and *WOX5* expression. We propose feed back loops to indicate *WOX5* induced cytokinin response, *NIN* expression and deconjugation mediated auxin response. The blunt arrows indicate Sugar and *CRE1* mediated cytokinin response to be mutually antagonistic. Dashed arrow indicates differential activity of *AhWOX5*.

Several evidences indicate a connection between sucrose signaling, the conserved WOX-CLV regulatory system, and the phytohormonal network that are involved in meristem maintenance (Francis and Halford, 2006). STIMPY (STIP; WOX9), a homeodomain transcription factor promotes WUS expression in the shoot meristem (Wu et al., 2005). Cytokinin signaling is required for *AtWOX9* expression (Skylar et al., 2010) and sucrose treatment could rescue *stip (AtWOX9)* mutants in *Arabidopsis* which has defects in SAM illustrating a cytokinin-WOX-sucrose module working in concert to control complex developmental processes (Wu et al., 2005). Accordingly in cytokinin receptor mutants, ectopic expression of *AtWOX9* could also restore establishment of SAM (Skylar et al., 2010). Similar example is restoration of shoot regenerative capacity in the type-B response regulator (*arr1* and *arr12*) mutants by ectopic expression of WUS (Meng et al., 2017). Overexpression of Fantastic 4 (*AtFAF4*) that controls meristem size by repressing WUS expression causes root growth arrest after germination and this aberration can also be rescued by sucrose treatment (Wahl et al., 2010). Our results further exemplify such sugar-WOX-phytohormone signaling axis by demonstrating turanose and WOX5 dependent recovery of RNS in a cytokinin perception mutant *cre1*. These examples point to a deep conservation of signaling for developing and sustenance of meristems with a direct link between WOX, sugar and phytohormonal networks (Gonzali et al., 2005; Wu et al., 2005; Wahl et al., 2010).

The homeobox transcription factors of WOX family are common regulators of cell proliferation and differentiation (Richards et al., 2015). The common mode of action for WOX5 proteins is prevention of premature differentiation by transcriptional repression and thereby maintaining a stem cell niche in meristems(Sarkar et al., 2007). The complete rescue of symbiosis in *cre1* by *AhWOX5* but not the intrinsic *MtWOX5* was intriguing because the reverse was actually expected (Fig.3; Supplementary table 2). This difference is possibly due to variation in post transcriptional or post translational regulations because (i) the transcript levels of both *WOX5* was similar in respective OE roots (Fig. 3E-F) and (ii) there was a significant overlap in the downstream responses induced by both the WOX5. Overexpression of WOX5 (*Mt* or *Ah*) had no effect on nodulation efficiency in wildtype background (A17) which further confirms WOX5 to be tightly controlled at post transcriptional level (Fig.3C; Supplementary Table 2). The homeobox domain is highly conserved between *Mt*WOX5 and *Ah*WOX5 with 96% similarity but there are notable features in AhWOX5 that distinguishes it from MtWOX5 for example (i) AhWOX5 has a 23 amino acid leader sequence in the N-terminus which is not present in MtWOX5 (Supplementary Fig. 3). This is important because intercellular mobility of WOX5 is important for stem-cell maintenance activity in roots (Pi et al., 2015) (ii) The WUS box motif is essential for WOX5 to act as transcriptional activator or repressor depending on developmental context and is absolutely essential for stem cell maintenance and interaction with TPL/TPR proteins (Leibfried et al., 2005; Ikeda et al., 2009; Dolzblasz et al., 2016). This motif is different between AhWOX5 (TL**Q**LFP) and MtWOX5 (TL**E**LFP). Earlier reports indicate substitutions in WUS box can completely alter the function of WOX proteins (Lin et al., 2013) (iii) AhWOX5 and MtWOX5 could be differentially regulated by the non-conserved regions in WOX proteins, as its essential role in determining the mobility and stability was previously noted in *Arabidopsis* (Daum et al., 2014). At present it is not clear which of these features of AhWOX5 are actually critical in the recovery of symbiosis in *cre1*, but the difference between MtWOX5 and AhWOX5 could emerge to be a model for understanding WOX5 action during nodule organogenesis in *M. truncatula*.

Nodule organogenesis in response to rhizobia is dependent on the cytokinin receptor CRE1 and the cytokinin-inducible transcription factor NIN (Gonzalez-Rizzo et al., 2006; Marsh et al., 2007; Plet et al., 2011; Schiessl et al., 2019). Interestingly, though ectopic expression of *MtWOX5* failed to restore symbiosis in *cre1*, it was functionally equivalent to *AhWOX5* in inducing *MtNIN* in both uninfected and infected roots of A17 and *cre1* (Fig.5A). This MtCRE1 and rhizobia independent mode of induction of *MtNIN* has not been noted earlier and highlights another layer to NIN regulation apart from the classical cytokinin-CRE1-NIN module that governs rhizobial symbiosis (Fig.6) (Laffont et al., 2020). Involvement of the paralogs *MtHK2/HK3* may be an unlikely option since WOX5 (*Mt* and *Ah*) expression has no effect on their expression (Fig.4B-C). Rather WOX5 might be directly targeting the response regulators downstream to CRE1 as previously observed in *Arabidopsis* SAM (Skylar and Wu, 2011). *MtNIN* expression also increased significantly by turanose treatment in both A17 and *cre1*, but only in infected roots (Fig.5A). Thus, turanose dependent *MtWOX5* is unable to trigger *MtNIN* expression in absence of rhizobia, which is unlike OE-WOX5 (*Mt* and *Ah*) roots where *MtNIN* expression increased in uninfected roots as well. Interestingly, we noted a significant inhibition of *MtCRE1* expression in turanose treated A17 roots and therefore turanose mediated *MtNIN* expression was MtCRE1 independent but absolutely dependent on upstream activation of SYM Pathway. These complex responses of *MtNIN* expression are in tune to distinct *cis*-regulatory sequences in NIN promoter that guide its complex spatio-temporal expression during symbiosis (Liu et al., 2019). Finally, just as turanose inhibited *MtCRE1* expression, the *pTCS:GUS* expression was always higher in turanose treated *cre1* highlighting a reciprocity in sugar and *MtCRE1* signaling (Fig.2G-H). Our results therefore suggest an alternate MtCRE1 independent module for *MtNIN* expression that is mediated through sugar and WOX5 and that reciprocally regulate the classical cytokinin-CRE1-NIN module (Fig. 6). Similar reciprocity between cytokinin and sugar signaling is noted earlier where plants with impaired cytokinin receptors display increased sugar sensitivity (Moore et al., 2003) and plants with decreased sensitivity to sugar was accompanied with increased cytokinin response (Franco-Zorrilla et al., 2005), suggesting a conserved relationship between sugar and cytokinin across plant species.

Both cytokinin and auxin responses are dependent on NIN action during symbiosis (Heckmann et al., 2011; Plet et al., 2011; Ng et al., 2015; Schiessl et al., 2019). Interestingly, though OE-*MtWOX5* and OE-*AhWOX5* was functionally equivalent in inducing *MtNIN*, there were distinct differences in the cytokinin and auxin responses that explains the difference in their ability to restore symbiosis in *cre1*. For example, the contrast in expression of *MtRR4*, a type-A response regulator could be critical. During symbiosis *MtRR4* expression depends on NIN and is associated with cortical cell division during nodule meristem formation (Vernié et al., 2015). In both OE-*AhWOX5* and OE-*MtWOX5*, expression of *MtRR4* increased in uninfected roots of A17 and *cre1* indicating a CRE1 independent pathway to induce the expression. In infected OE-*AhWOX5* roots expression of *MtRR4* further increased (Fig.4D). But in infected roots of OE-*MtWOX5*, the expression of *MtRR4* significantly decreased specifically in *cre1* roots but not in A17 indicating MtCRE1 signaling to be essential for *MtRR4* expression in infected roots. Earlier reports have shown WOX proteins to directly regulate the expression of Type-A RRs genes (Zhao et al., 2009) and the distinct differences between AhWOX5 and MtWOX5 could be the reason behind the differences in *MtRR4* expression.

Auxin accumulation (*pDR5:GUS* expression) depends on NIN, as it does not occur in a *nin* null mutant (Suzaki et al., 2012). Downstream to NIN, the genes that regulate auxin availability was differentially affected by WOX5 (*Ah* and *Mt*) and turanose (Fig.5). The most distinctive was the expression of an auxin conjugate hydrolase *MtIAR33* that significantly increased in OE-*AhWOX5* and turanose treated roots indicating a WOX5 dependent pathway to upregulate *MtIAR33*. Expression of *MtIAR33* in OE-*AhWOX5* roots or turanose treated roots was equivalent or more in *cre1* as compared to A17 indicating a CRE1 independent pathway to induce *MtIAR33* expression (Fig.5E). Again auxin response (*pDR5:GUS*) in turanose treated *cre1* roots was significantly higher than auxin response in A17 roots indicating CRE1 restricts sugar mediated auxin response (Fig.2K and O). Taken together these results prompt us to hypothesize that the turanose-WOX5 signaling axis primarily functioned through *MtNIN* and *MtIAR33*, and hydrolysis of auxin conjugates was preferentially used by this axis in determining the bioactive pool of auxin for sugar mediated restoration of symbiosis in *cre1*. There was no change in expression of *MtIAR33* in OE-*MtWOX5* indicating the importance of deconjugation of auxin in restoration of symbiosis in *cre1*. Thus, apart from auxin transport (Rightmyer and Long, 2011; Ng et al., 2015), and biosynthesis (Schiessl et al., 2019), that functions in an MtCRE1 dependent manner, our results suggest a CRE1 independent auxin deconjugation to be a potential pathway for auxin accumulation.

Several earlier evidences indicate auxin responsive WOX5 expression (Ding and Friml, 2010; Osipova et al., 2012) whereas in our results WOX5 dependent IAR33 expression suggest auxin accumulation to happen downstream to WOX5. This is similar to a previous report where turanose treatment inhibited auxin conjugation to increase free auxin content but failed to do so in turanose insensitive (*tin*) loss of function *wox5* mutants thereby indicating WOX5 to mediate the accumulation of bioactive auxin pool (Gonzali et al., 2005). It is possible that WOX5 mediates a positive feedback loop for rapid auxin response (Fig.6). We envisage this positive loop to involve NIN because a WOX-NIN crosstalk would mean a cross talk between a sugar sensing and a nitrogen sensing network being connected during inception of symbiosis. It may be noted that *MtIAR33* was significantly upregulated during early stages of rhizobial infection in *M. truncatula* (Campanella et al., 2008). However, there was no upregulation of *MtIAR33* in our experimental conditions in absence of turanose and the difference is most likely due to presence of β-aminobutyric acid (BABA) in their nodulation media that is known for having priming effects on plants against stress responses (Rejeb et al., 2018). In analogy, it is likely that turanose treatment and *AhWOX5* expression actually primed the *cre1* roots with increased expression of *MtIAR33* that developed a CRE1 independent robust auxin response and thereby restored symbiosis in *cre1*. Incidence of merged nodules which is an indication of higher auxin concentration was only observed in presence of turanose which could be due to increased expression of *MtIAR33* in presence of turanose or a better homeostatic control over auxin availability in presence of *AhWOX5* (Penmetsa et al., 2003; van Noorden et al., 2007; Suzaki et al., 2012). For example, OE-*AhWOX5*, but not the turanose treated roots have increased expression of *MtLBD16* and *MtYUCCA8* and might therefore have better control over auxin response that led to higher efficacy of restoration of symbiosis in *cre1* than turanose treated plants that recruited only auxin deconjugation (*MtIAR33*).

In a working model we propose how turanose-wox5 signaling might have restored symbiosis in *cre1* (black arrow) in the backdrop of the present understanding about cytokinin signaling (grey arrows) during nodule initiation in *M. truncatula* (Fig.6). We propose a turanose dependent WOX5 expression to turn on feed forward positive loop(s) to increase cytokinin and auxin response and NIN expression. This turanose-WOX5 signaling axis is CRE1 independent and reciprocally regulates the classical cytokinin-CRE1-NIN module. Future work is needed to understand the basis of differential action of MtWOX5 and AhWOX5 and the role of NIN in deconjugation mediated auxin accumulation. We also need to better understand the sugar perception and signaling network to understand its role in nodule meristem development.

## MATERIALS AND METHODS

### Plant and rhizobial strains

*Medicago cre1* seeds (Plet et al., 2011), *Agrobacterium rhizogenes* strain MSU440 and *Sinorhizobium meliloti Sm2011-pBHR-mRFP* and *Sm1021-pXLGD4-lacZ* (Boivin et al., 1990) strains were used.

### Medicago growth conditions

*Medicago* was grown as described in (Garcia et al., 2006). In brief, seeds were scarified for 10min in H_2_SO_4_, rinsed with sterile water, followed by surface sterilization with 6% sodium hypochlorite for 3 min containing a few drops of Tween-20 and then again rinsed with sterile water. Surface-sterilized seeds were imbibed with sterile water with gentle rotation for 5 to 7h at ambient temperature on a rotating shaker and then stored in water at 4° C overnight. Seeds were then placed on inverted plates containing moist filter papers and then left in dark at 20^°^Cfor 36 h for germination. Germinated seedlings were placed in Fahraeus (Catoira et al., 2000) agar (0.8%) medium in square petri dishes (12 × 12 cm) and maintained in a growth chamberat 20°C on a 16-2 h-light and 8-h-dark schedule under 100-110 μmol/m /s.

### Turanose and sucrose treatment

Plants are germinated as previously described by (Saha et al., 2014) followed by their transfer into Fahraeus medium containing different concentration of turanose and sucrose:10^−4^M, 10^−3^M and 10^−2^M for 1 week. For promoter assay plants are maintained for 1 week in Turanose plates before infected with *Sm2011-pBHR-mRFP* for 24 hours then plants are harvested and stained. For nodulation assay plants are inoculated in the plates with *Sm2011-pBHR-mRFP/ Sm1021-pXLGD4-lacZ* grown overnight at 28°C in YM to OD600=1.0 and then diluted 1:50 in half-strength B&D (Broughton and Dilworth, 1971). From this diluted bacterial culture 2ml/plate was used for inoculation. Nodulation is scored 3WAI.

### Constructs

Full length MtWOX5 and AhWOX5 are amplified from cDNA prepared from nodulated roots of *Medicago truncatula* and *Arachis hypogaea* (Kundu and DasGupta, 2018) with primers 5’-CACCATGGAAGAGAGCATGTCAGG-3’ and 5’ - ACTTACGGTTGAGTTTTGTGTAA-3’, and (*MtWOX5*) 5’-CACCATGCAGACGGTCCGAGATCTGTC-3’ and 5’- CCTTCGCTTAAGTTTCATGTAA-3’ (*AhWOX5*) and cloned into pENTR-dTOPO (Invitrogen). The entry clones of *AhWOX5 & MtWOX5* recombined through LR clonase Gateway technology (Invitrogen) into *pK7WGF2* to generate *p35S::eGFP-AhWOX5 p35S::eGFP-MtWOX5* respectively. *pDR5:GUS, pWOX5:GUS* and *pTCS:GUS* were obtained from (Franssen et al., 2015).

### Induction of hairy roots and generation of composite *M. truncatula* plants and scoring nodulation

Seedlings were transformed with indicated constructs in *Agrobacterium rhizogenes* MSU440 by following a standard procedure (Boisson-Dernier et al., 2001). Briefly, when seedlings had a radicle length of approximately 10 mm, the radicle was sectioned approximately 3 mm from the root tip with a sterile scalpel. Sectioned radicles were inoculated by coating the freshly cutsurface with *A. rhizogenes* grown on TY solid medium. Thereafter, the inoculated sectioned seedlings were placed on slanted agar (Sigma) in square petri dishes containing Fahraeus medium with kanamycin at 25 mg/liter and maintained at 20°C. Within 3-4 weeks, transgenic hairy roots were screened for GFP fluorescence by Leica stereo fluorescence microscope M205FA equipped with a Leica DFC310FX digital camera (Leica Microsystems). The selected composite transgenic plants were transplanted in vermiculite pots and the plants were maintained in a growth chamber at 2 20°C on a 16-h-lightand 8-h-dark schedule under 100-110 μmols/m /s. For nodulation assay plants are inoculated with *Sm2011-pBHR-mRFP/ Sm1021-pXLGD4-lacZ* grown overnight at 28°C in YM to OD600=1.0 and then diluted 1:50 in half-strength B&D (Broughton and Dilworth, 1971). From this diluted bacterial culture 2ml/pot was used for inoculation. Nodulation was scored 4 and 6WAI with *S. meliloti* expressing mRFP (Smit et al., 2005). For observing ITs, plants were harvested 2WAI.

### Acetylene reduction assay

The nitrogen-fixing activity of the nodules was examined indirectly by Acetylene Reduction Assay as described by (Koch and Evans, 1966) using gas chromatography, the bacterial nitrogenase activity of the nodules at 6WAI with *Sm2011-pBHR-mRFP* was measured. The roots were placed into 10mL glass vials sealed with rubber septa, and 200μL of acetylene was injected into each vial, incubated for 90 min at room temperature, and ethylene production was measured using Clarus 460 PerkinElmer Gas Chromatograph. The total nitrogenase activity was calculated as ARA units (nanomoles of ethylene. Hour^−1^ mg nodule^−1^). Three independent biological replicates were performed, with n>4 plants analyzed per replicate. Nodules from individual plant root systems were then pulled out and weighted to determine specific nitrogenase activity (ARA units per mg of nodule).

### Phenotypic analysis, histochemical staining and confocal microscopy

Images of whole mount nodulated roots were captured using a Leica stereo fluorescence microscope M205FA equipped with a Leica DFC310FX digital camera (Leica Microsystems). Histological assay for checking GUS expression was performed according to Saha et. al.,2014 with a short staining period. Briefly, roots were washed with 0.1 M potassium phosphate buffer (pH 7.0) and immersed and incubated in the dark in staining solution 1 mM 5-bromo-4-chloro-3-indolyl-β-D-glucuronicacid (X-Gluc, Thermo Scientific), 50mM sodium phosphate buffer, 0.5mM potassium ferrocyanide, 0.5mM potassium ferricyanide, 20% methanol and 0.1%Triton X-100 for 15 minutes at 37°C. After rinsing in phosphate buffer and cleaning in 70% ethanol, the roots were analyzed and imaged using Leica stereo fluorescence microscope M205FA (Leica Microsystems) and Olympus fluorescence microscope (IX71, Japan). Sectioning of Gus stained roots was done using cryo microtome (Leica Microsystem) and sliced into 40-μm thick sections and imaged using Olympus fluorescence microscope (IX71, Japan). Detached nodules were embedded in 5% low melting point agarose and sliced into 30-μm thick sections with a rotary microtome RM2235 (Leica Microsystems). Roots were also stained with X-Gal (Affymetrix) for lacZ-activity, asdescribed in (Arrighi et al., 2008) and imaged under an Olympus IX71 microscope (Japan). For confocal microscopy, sample preparation was done according to Haynes and associates (Haynes et al., 2004). Images were acquired with a Leica TCS SP5 II AOBS confocal laser scanning microscope (Leica Microsystems). All digital micrographs wereprocessed using Adobe Photoshop CS6.

### Real time PCR

Superscript III RT (Life Technologies) and anchored dT17 primers as described in the manufacturer’s protocol. 1μl of cDNA was used to set up real time qRT-PCR in a 20µl reaction system with 2X SYBR Green PCR master mix (Life Technologies). Experimental setup and execution were conducted using an ABI 7500 Fast Real-Time PCR system. PCR program: 1cycle at 50°C, 1cycle at 95°C for 5 min, 40 cycles at 95°C for 30s and 52°C for 30s. Expression data were obtained from three independent biological repetitions. Calculations were done using the cycle threshold method using *MtACTIN2* as the endogenous control. All primers including the genes used for normalization are given in (Supplementary Table 3). Statistical significance was determined based on Mann-Whitney test.

### Accession number

Nucleotide sequence data for the genes used in thisarticle can be found in the GenBank data library under the following accession numbers: MtCRE1, XM_024773926.1;*Mt*HK2, XP_003617960.1; *Mt*HK3, XP_003601762.1; MtRR4, XM_003613375.3; MtRR8, 003608889.3; MtRR9, XM_024778303.1;*Mt*NIN, FJ719774; MtLBD16, XM_013594406.2; MtYUCCA2, XM_013597647.2; MtYUCCA8, XM_003625408; MtIAR33, DQ489994; MtPIN4, AY115839.1;*Mt*ACTIN2, JQ02873.

## ACKNOWLEDGEMENT

We thank Florian Frugier for *cre1* seeds; Henk J. Franssen for providing us with *pDR5:GUS, pTCS:GUS* and *pWOX5:GUS* constructs; Ton Bisseling & Erik Limpens for *S. meliloti* harboring *pBHR-mRFP;* Douglas R. Cook for *Agrobacterium* strain MSU440.

## SUPPLEMENTAL DATA

**Supplementary Table 1:** Nodule number in A17 and *cre1* upon turanose and sucrose treatment.

**Supplementary Table 2:** Nodule number in A17and *cre1* by overexpression of *AhWOX5* and *MtWOX5*.

**Supplementary Table 3:** List of primers used for cloning and qRT-PCR.

**Supplementary figure 1:** Effect of Turanose and Sucrose treatment on Nodulation in A17 and *cre1*.

**Supplementary figure 2:** *MtWOX5* expression in response to turanose treatment.

**Supplementary figure 3:** Sequence alignment and phylogenetic tree of WOX5.

**Supplementary figure 4:** IT morphology and nodule number under WOX5 overexpression.

**Supplementary figure 5:** Relative expression of *MtWOX5* and *AhWOX5* in A17.

## Notes

**Financial support:** This work was funded by Grants from Govt. of India: DST (DST EMR/2015/001006). Fellowship to A. K (Council of Scientific and Industrial Research, CSIR-09/028[0756]/2009– EMR–I) and fellowship to F.M (University Grant Commission, No.F.16-6(DEC.2016)/2017 (NET). UGC Ref no- 771

The authors declare no conflict of interest.

### Competing Interest Statement

The authors have declared no competing interest.

### Summary of Updates

Results and Discussion is revised in this manuscript version emphasizing the role of MtIAR33 in the rescue of nodulation in cre1

