## Supplementary figures and tables for "Turanose induced WOX5 restores symbiosis in the *Medicago truncatula* cytokinin perception mutant *cre1*"

**Supplementary Table 1:** Nodule number in A17 and *cre1* upon turanose and sucrose treatment.

**Supplementary Table 2:** Nodule number in *cre1* and A17 of *M. truncatula* by overexpression of *AhWOX5* and *MtWOX5*.

**Supplementary Table 3:** List of primers used for cloning and qRT-PCR.

**Supplementary figure 1:** Effect of Turanose and Sucrose treatment on Nodulation in A17 and *cre1*.

**Supplementary figure 2:** *MtWOX5* expression in response to turanose treatment.

**Supplementary figure 3:** Sequence alignment and phylogenetic tree of WOX5.

**Supplementary figure 4:** IT morphology and nodule number under WOX5 overexpression.

**Supplementary figure 5:** Relative expression of *MtWOX5* and *AhWOX5* in A17.

**Supplementary table 1:**

**Nodule number in A17 and *cre1* upon turanose and sucrose treatment.**

| **Treatment** | **Nodule number in A17** | | **Nodule number in *cre1*** | |
|---|---|---|---|---|
| | **white nodules** | **pink nodules** | **white nodules** | **pink nodules** |
| Control | 2.2 (N=10/10) | 18.6 (N=10/10) | 1 (N=4/10) | 0.41 (N=4/10) |
| $10^{-4}$M Turanose | 3.5 (N=20/20) | 16.4 (N=20/20) | 5.4 (N=16/20) | 2.2 (N=16/20) |
| $10^{-3}$M Turanose | 11.6 (N=25/25) | 17.6 (N=25/25) | 7.16 (N=18/20) | 3.25 (N=18/20) |
| $10^{-2}$M Turanose | 5.8 (N=20/20) | 16.9 (N=20/20) | 6.9 (N=17/20) | 2.9 (N=17/20) |
| $10^{-4}$M Sucrose | 1.55 (N=9/9) | 10.55 (N=9/9) | 0.33 (N=9/12) | 1.08 (N=9/12) |
| $10^{-3}$M Sucrose | 0.7 (N=9/10) | 2.1 (9/10) | 0.41 (N=8/12) | 0.66 (N=8/12) |
| $10^{-2}$M Sucrose | 0.6 (N=6/10) | 0.5 (N=6/10) | 0.5 (N=6/12) | 0.16 (N=6/12) |

N=Nodulated plants/plants studied

Nodule number (Pink and White) in A17 (WT) and *cre1* (cytokinin perception mutant of *Medicago truncatula*) by turanose and sucrose treatment 3WAI with *Sm2011-pBHR-mRFP*

**Supplementary table 2:**

**Nodule number in *cre1* and A17 of *M. truncatula* by overexpression of *AhWOX5* and *MtWOX5*.**

| Genotype | Construct | Nodules/Plant | |
|---|---|---|---|
| A17 | | **white nodules** | **pink nodules** |
| | Control vector (ev) | NA (N=5/5) | 43 ± 12.4 (N=5/5) |
| | 35S:*MtWOX5*(N-Terminal gfp) | NA (N=5/5) | 40 ± 10.0 (N=5/5) |
| | 35S:*AhWOX5*(N- Terminal gfp) | NA (N=5/5) | 43 ± 10.2 (N=5/5) |
| *cre1* | Control vector (ev) | 1.6 ± 1.0 (N=2/5) | 0.6 ± 0.8 (N=2/5) |
| | 35S:*MtWOX5*(N-Terminal gfp) | 4 ± 4.2 (N=4/10) | 2.1 ± 2.6 (N=4/10) |
| | 35S:*AhWOX5*(N- Terminal gfp) | 19.33 ± 4.8 (N=24/25) | 46 ± 15.6 (N=24/25) |

N=Nodulated plants/plants studied.

Nodule number (Pink and White) in A17 (WT) and *cre1* (cytokinin perception mutant) by overexpression of Control vector (ev), *p35S:MtWOX5* (N-Terminal gfp), *p35S:MtWOX5* (C-Terminal gfp), *p35S:AhWOX5* (N- Terminal gfp) in plants grown in vermiculite.

**Supplementary table 3: List of primers used for cloning and qRT-PCR.**

| Primer name | 5'-3' Sequences |
|---|---|
| MtCRE1 qFor | GTCGCCGCCGGTGCATTAAAGAAG |
| MtCRE1 qRev | CCCGCACTTCAAGCACTCTTC |
| MtHK2 qFor | CAGCTTCATGTTTTGGCTTCCTTGTTC |
| MtHK2 qRev | GCAGAGACCAATGTTACATATGCTTTCC |
| MtHK3 qFor | CCATGGAGCAGAACCCGG |
| MtHK3 qRev | CTCTCTTGTAAACAGCAAATGTGAGG |
| MtRR4 qFor | GTTCCGGGTTTAAAGGTGGATCT |
| MtRR4 qRev | GACGTGAGATAATTTCACTGGCT |
| MtRR8 qFor | GATTTCTTCCTGCAAAGTGACGGTTG |
| MtRR8 qRev | CCTCTGCTCCTTCCTCCAAACACC |
| MtRR9 qFor | CATGTGTTGGCTGTTGATGACAG |
| MtRR9 qRev | GCAAATCATAGCCAGTCATCCCAGGC |
| MtNINqFor | GATCGTCACCATCCAAGAAAACCCAC |
| MtNINqRev | CTGCTGCTGATGAAACTATTATGCTTTGTATCC |
| MtLBD16 qFor | AGCTCGTATCAGAGACCCTGT |
| MtLBD16 qREV | GACTCAAGTGAGCTTTGAGGAG |
| MtYUCCA2 qFor | GGGTGTGGAAATTCAGGTATGGAG |
| MtYUCCA2 qRev | CCACAAATTGAACAGAGAACC |
| MtYUCCA8 qFOR | GCTAACAAATTTGAAATCAACCCG |
| MtYUCCA8 qRev | GTCCTTCAATTTCAGGAGTAACAC |
| MtIAR33 qFor | GCTATTCGTGCTGATATCGATGGAC |
| MtIAR33 qRev | CCCTCCTCAGCAGGTTGGAAAAG |
| MtPIN4 qFor | GCATGGCTATGTTCAGTCTTGG |
| MtPIN4 qRev | GACCAACAAGGAATCTCACACC |
| MtWOX5 qFor | GGCACAAAGTGTGGTCGTTGGAATCC |
| MtWOX5 qRev | GAAACCAATAGAACACATTCTTGC |
| MtWOX5 For | CACCATGGAAGAGAGCATGTCAGG |
| MtWOX5 Rev | ACTTACGGTTGAGTTTTGTGTAA |
| AhWOX5 qFor | GGAACAAAGTGTGGGCGTTGG |
| AhWOX5 qRev | CTCTCCCTAGCCTTATGATTCTGAAACC |
| AhWOX5 For | CACCATGCAGACGGTCCGAGATCTGTC |
| AhWOX5 Rev | CCTTCGCTTAAGTTTCATGTAA |
| MtActin2 qFor | ATGGAGAAGATCTGGCATCA |
| MtActin2 qRev | ATACCTGTTGTACCGACCACT |

**SUPPLEMENTARY FIGURE LEGENDS:**

**Supplementary figure 1: Effect of Turanose and Sucrose treatment on Nodulation in A17 and *cre1*.**Roots were treated with$10^{-4}$M to$10^{-2}$ M Turanose and sucrose for 7 days and infected with *Sm2011-pBHR-mRFP.* At 3 WAI(A)Root length, (B) Shootlength, (C) Number oflateral roots/plant and (D) Nodule number/plant was recorded. Histogram represents anaverage ofthree biological replicates each having n>6 plants and error bar represents SD.Mann-Whitney test wasused to assess significant differences, where ***, **and *indicate P<0.0007, 0.006 and 0.05 respectively.

**Supplementary figure 2: *MtWOX5* expression in response to turanose treatment.** A17 and *cre1* roots were treated with $10^{-3}$M Turanose for 7 days before infection with *Sm-2011-pBHR-mRFP*. qRT-PCR analysis of *MtWOX5* in turanose treated (infected and uninfected) A17 and *cre1* roots are shown relative to uninfected/untreated Al7 roots. Histogram represents anaverage of three biological replicates each having n>4 plants anderror bar represents SD. Mann-Whitney test was used to assess significant differences, where **indicate P<0.005.

**Supplementary figure 3: Sequence alignment and phylogenetic tree of legume WOX5.** (A)CLUSTALW alignment of WOX5 protein from *Glycine max* (XP_003537483), *Vigna angularis* (XP_017416822), *Cajanus cajan* (XP_020202518), *Lotus japonicus* (Lj0g3v0135189), *Arachis hypogaea* (KT820790), *Pisum sativum* (AEX88468.1), *Medicago truncatula* (XP_003616581.1), *Trifoliumsubterraneum* (GAU36366.1), *Cicer arietinum* (XP_012568542.1), *Nicotiana tabacum* (XP_016445580.1) and *Arabidopsis thaliana* (NP_187735.2) where the boxes represents Homeodomain, acidic domain, WUS domain and EAR domain. Amino acids are coloured as conserved (red), partially conserved (blue) and non-conserved (black). Solid bars below the homeodomain represent the conserved helix I, II and III (Lian et al., 2014) and arrowhead represent the difference in amino acid in the conserved WUS domain among the legumes. (B) Neighbour-joining distance tree (Saitou and Nei, 1987) of WOX5s using MEGA6 (Tamura et al., 2013) based on CLUSTAL W alignment of the amino acids where cyan and green branches represent indeterminate and determinate nodulators respectively.

**Supplementary figure 4: IT morphology and nodule number under WOX5 overexpression.** Epidermal infection thread (IT) observed 2WAI with *Sm1021-pXLGD4-lacZ* in *cre1* roots transformed with **(A-C)** empty vector, **(D-F)***p35S::eGFP-MtWOX5* and **(G-I)***p35S::eGFP-AhWOX5*. Each of the infection events has been roughly classified into 4 categories: (i) successful cortical invasion and intracellular colonization (G-I) (ii) Stalled in nodule apex (C and F), (iii) stalled in epidermal cortical barrier (B and E) and (iv) stalled in microcolonies in root hair (A and D). Category (i) is considered normal whereas Category (ii-iv) is considered as abnormality. **(J)** Box plot represents total nodule number per plant in empty vector, *p35S::eGFP-MtWOX5* and *p35S::eGFP-AhWOX5* transformed hairy-root systems in A17 and *cre1* respectively, harvested 4 weeks after infection (4WAI) with *Sinorhizobium meliloti Sm2011-pBHR-mRFP*. Mann-Whitney test was used to assess significant differences, where **** and *** indicates P<0.0001 and 0.0002 respectively.

**Supplementary figure 5: Relative expression of *MtWOX5* and *AhWOX5* in A17. (A-B)** qRT-PCR analysis of (A) *MtWOX5* and (B) *AhWOX5* relative to empty vector transformed (A17) roots normalized against *MtActin*.(A-B)Histogram represent an average of three biological replicates each having n>4 plants and error bar represents SD. Mann-Whitney test was used to assess significant differences, where* indicate P<0.02.

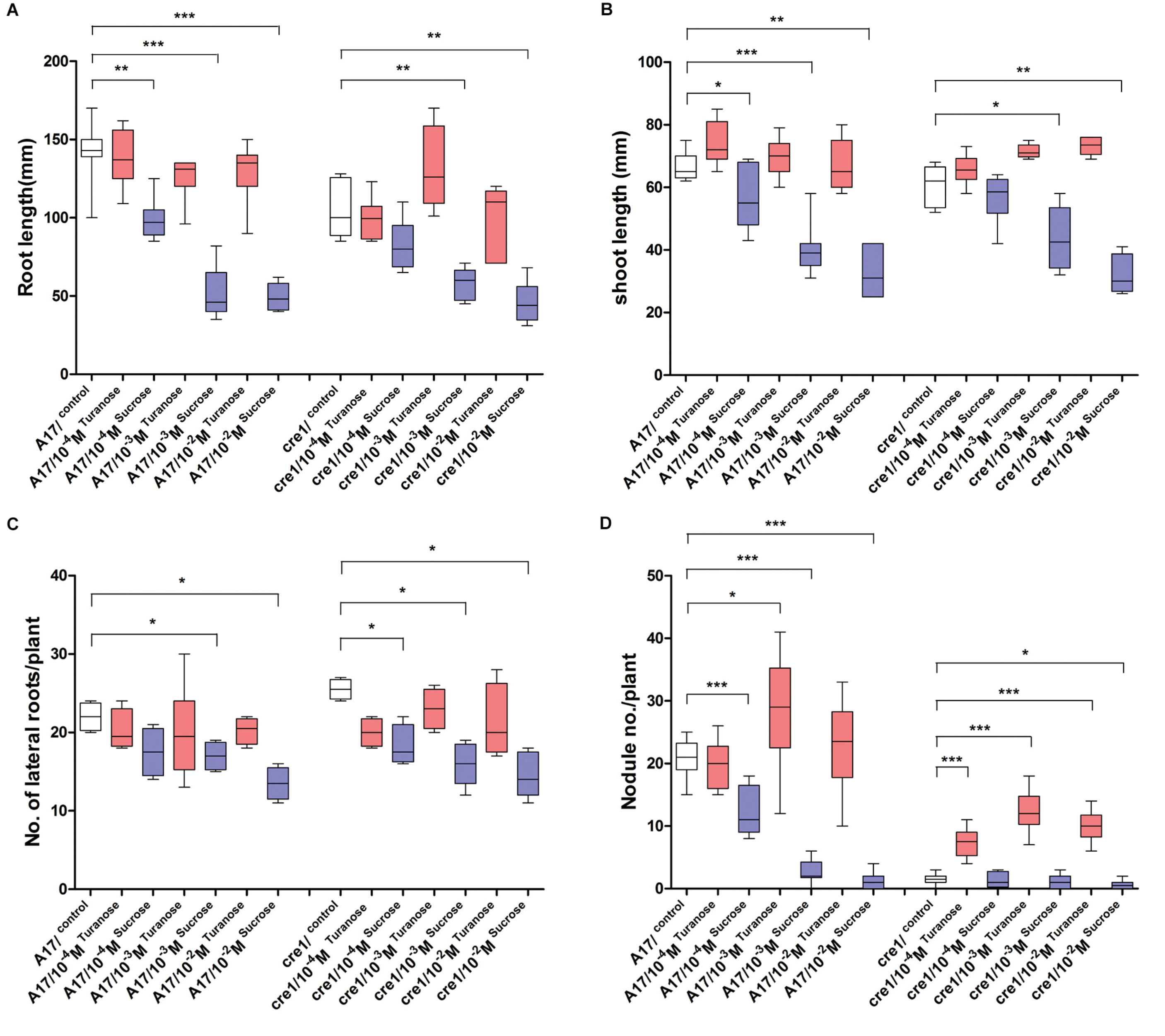


**Supplementary figure 1: Effect of Turanose and Sucrose treatment on Nodulation in A17 and *cre1*.** Roots were treated with $10^{-4}$M to $10^{-2}$ M Turanose and sucrose for 7 days and infected with *Sm2011-pBHR-mRFP*. At 3 WAI (A) Root length, (B) Shoot length, (C) Number of lateral roots/plant and (D) Nodule number/plant was recorded. Histogram represents an average of three biological replicates each having n>6 plants and error bar represents SD. Mann-Whitney test was used to assess significant differences, where ***, ** and * indicate P<0.0007, 0.006 and 0.05 respectively.

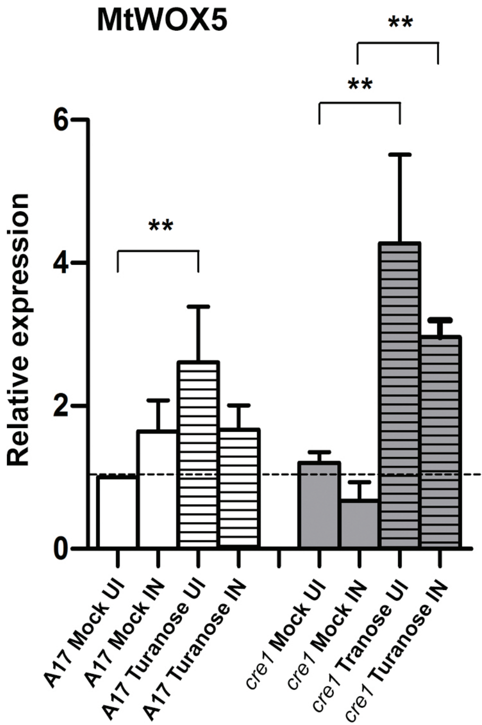


**Supplementary figure 2: *MtWOX5* expression in response to turanose treatment.** A17 and *cre1* roots were treated with 10 $^{-3}$M Turanose for 7 days before infection with *Sm-2011-pBHR-mRFP*. qRT-PCR analysis of *MtWOX5* in turanose treated (infected and uninfected) A17 and *cre1* roots are shown relative to uninfected/untreated Al7 roots. Histogram represents an average of three biological replicates each having n>4 plants and error bar represents SD. Mann-Whitney test was used to assess significant differences, where ** indicate P<0.005.

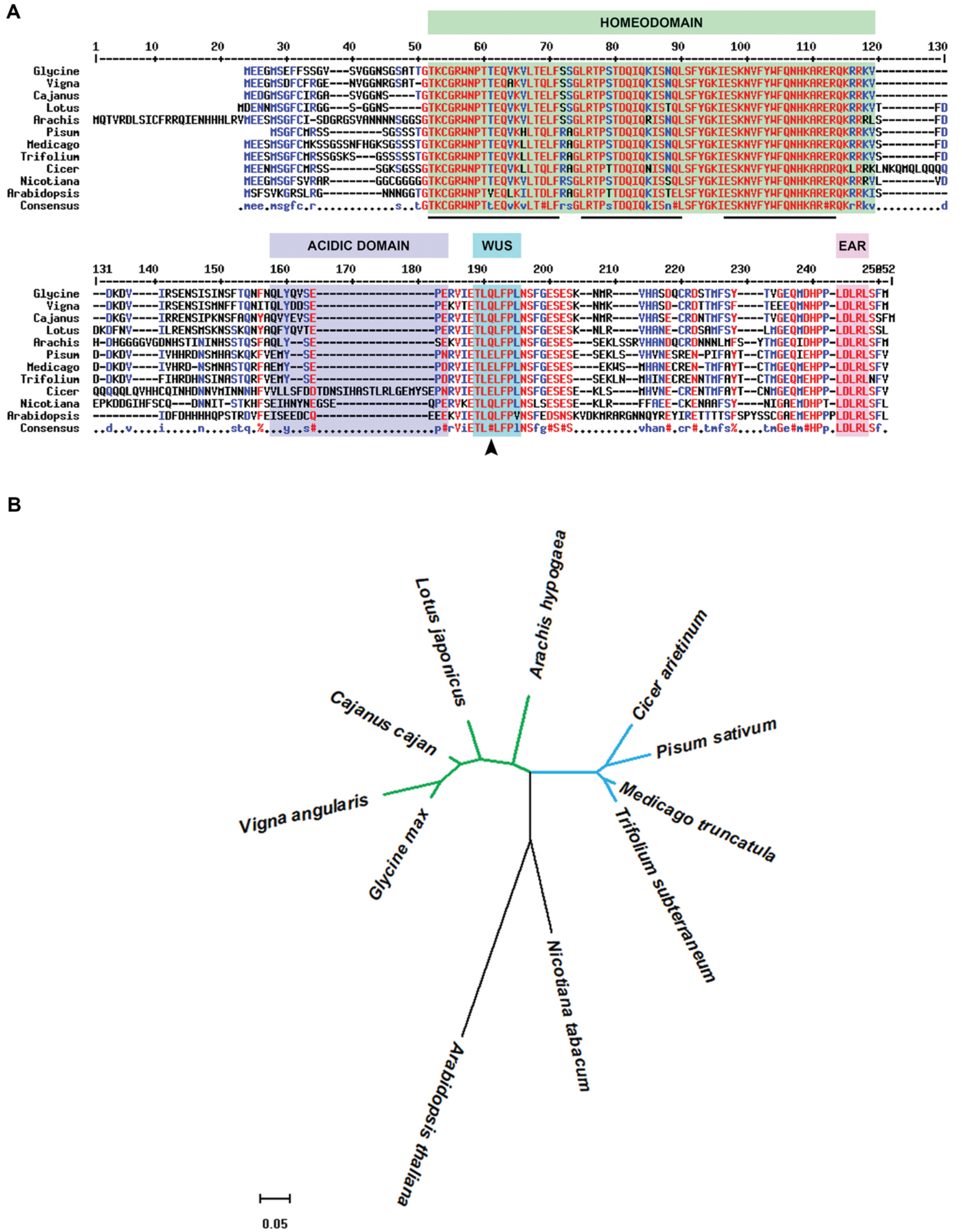


**Supplementary figure 3: Sequence alignment and phylogenetic tree of legume WOX5.** (A) CLUSTALW alignment of WOX5 protein from *Glycine max* (XP_003537483), *Vigna angularis* (XP_017416822), *Cajanus cajan* (XP_020202518), *Lotus japonicus* (Lj0g3v0135189), *Arachis hypogaea* (KT820790), *Pisum sativum* (AEX88468.1), *Medicago truncatula* (XP_003616581.1), *Trifolium subterraneum* (GAU36366.1), *Cicer arietinum* (XP_012568542.1), *Nicotiana tabacum* (XP_016445580.1) and *Arabidopsis thaliana* (NP_187735.2) where the boxes represents Homeodomain, acidic domain, WUS domain and EAR domain. Amino acids are coloured as conserved (red), partially conserved (blue) and non-conserved (black). Solid bars below the homeodomain represent the conserved helix I, II and III (Lian et al., 2014) and arrowhead represent the difference in amino acid in the conserved WUS domain among the legumes. (B) Neighbour-joining distance tree (Saitou and Nei, 1987) of WOX5s using MEGA6 (Tamura et al., 2013) based on CLUSTAL W alignment of the amino acids where cyan and green branches represent indeterminate and determinate nodulators respectively.

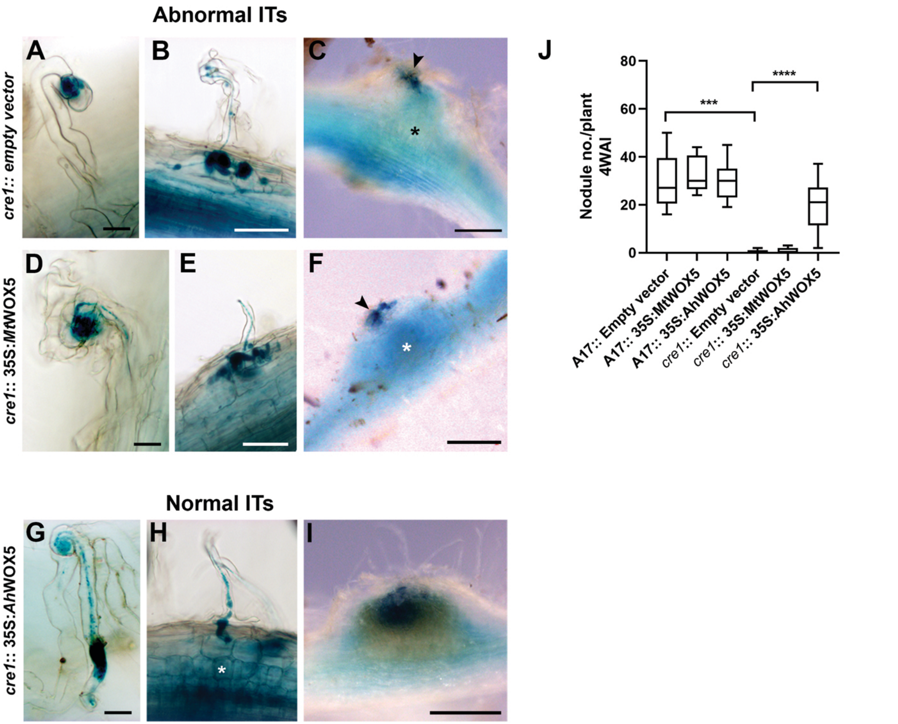


**Supplementary figure 4: IT morphology and nodule number under WOX5 overexpression.** Epidermal infection thread (IT) observed 2WAI with *Sm1021-pXLGD4-lacZ* in *cre1* roots transformed with **(A-C)** empty vector, **(D-F)** *p35S::eGFP-MtWOX5* and **(G-I)** *p35S::eGFP-AhWOX5*. Each of the infection events has been roughly classified into 4 categories: (i) successful cortical invasion and intracellular colonization (G-I) (ii) Stalled in nodule apex (C and F), (iii) stalled in epidermal cortical barrier (B and E) and (iv) stalled in microcolonies in root hair (A and D). Category (i) is considered normal whereas Category (ii-iv) is considered as abnormality. **(J)** Box plot represents total nodule number per plant in empty vector, *p35S::eGFP-MtWOX5* and *p35S::eGFP-AhWOX5* transformed hairy-root systems in A17 and *cre1* respectively, harvested 4 weeks after infection (4WAI) with *Sinorhizobium meliloti Sm2011-pBHR-mRFP*. Mann-Whitney test was used to assess significant differences, where **** and *** indicates $P<0.0001$ and 0.0002 respectively.

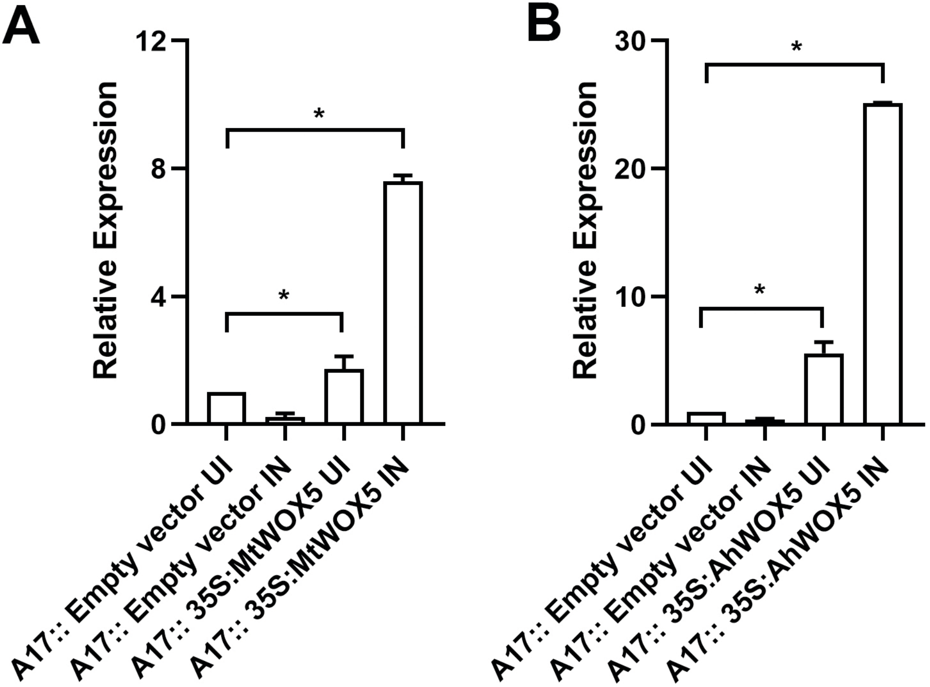


**Supplementary figure 5: Relative expression of *MtWOX5* and *AhWOX5* in A17. (A-B)** qRT-PCR analysis of (A) *MtWOX5* and (B) *AhWOX5* relative to empty vector transformed (A17) roots normalized against *MtActin*. (A-B) Histogram represent an average of three biological replicates each having n>4 plants and error bar represents SD. Mann-Whitney test was used to assess significant differences, where * indicate P<0.02.
